# Liver-innervating vagal sensory neurons are indispensable for the development of hepatic steatosis and anxiety-like behavior in diet-induced obese mice

**DOI:** 10.1101/2024.02.20.581228

**Authors:** Jiyeon Hwang, Sangbhin Lee, Junichi Okada, Li Liu, Jeffrey E. Pessin, Streamson C. Chua, Gary J. Schwartz, Young-Hwan Jo

**Affiliations:** The Fleischer Institute for Diabetes and Metabolism; Division of Endocrinology, Department of Medicine; Department of Molecular Pharmacology; Department of Neuroscience, Albert Einstein College of Medicine, NY, USA

**Author notes:** **Corresponding author:** Departments of Medicine, Molecular Pharmacology, and Neuroscience Albert Einstein College of Medicine, 1300 Morris Park Ave, Bronx, NY 10461.

## Abstract

The visceral organ-brain axis, mediated by vagal sensory neurons, is essential for maintaining various physiological functions. Here, we investigate the impact of liver-projecting vagal sensory neurons on energy balance, hepatic steatosis, and anxiety-like behavior in mice under obesogenic conditions. A small subset of vagal sensory neurons in both the left and right ganglia innervate the liver and project centrally to the nucleus of the tractus solitarius, area postrema, and dorsal motor nucleus of the vagus, and peripherally to the periportal areas in the liver. Surprisingly, the loss of liver-projecting vagal sensory neurons via caspase-induced selective destruction of advillin-positive neurons prevents diet-induced obesity, and these outcomes are associated with increased energy expenditure. Although males and females exhibit improved glucose homeostasis following disruption of liver-projecting vagal sensory neurons, only male mice display increased insulin sensitivity. Furthermore, the loss of liver-projecting vagal sensory neurons limits the progression of hepatic steatosis in mice fed a steatogenic diet. Intriguingly, mice lacking liver-innervating vagal sensory neurons also exhibit less anxiety-like behavior compared to control mice. Therefore, modulation of the liver-brain axis may aid in designing effective treatments for both psychiatric and metabolic disorders associated with obesity and MAFLD.

## Introduction

Communication between the visceral organs and the brain is required to maintain metabolic homeostasis. Vagal sensory neurons present in the vagus nerve ganglia transmit interoceptive signals from visceral organs to the medulla^1^. Vagal sensory neurons are highly heterogeneous in their molecular identities, anatomical connections, and physiological roles^1–3^. Each type of vagal sensory neuron with unique molecular features has a specific target organ such as the larynx, stomach, intestine, pancreas, heart, and lung^3–10^. The highly specialized cellular, molecular, and anatomical organization of vagal sensory neurons has been shown to be crucial for the proper detection and integration of interoceptive signals to control metabolic homeostasis and other vital functions, such as breathing and reward^1–10^.

The molecular identity of vagal sensory neurons appears to play a major role in determining their functional properties^3,5–8,10^. For example, vagal sensory neurons that control the digestive system include vasoactive intestinal peptide-, glucagon-like peptide 1 receptor-, oxytocin receptor-, and G protein-coupled receptor 65-expressing neurons^5,8^. Interestingly, distinct types of vagal sensory neurons (somatostatin- vs. CGRP-positive cells) innervate different parts of the stomach^5^. Those innervating the lungs are purinergic receptor P2RY1- positive neurons and neuropeptide Y receptor Y2-expressing neurons^6^. Pancreatic islets are innervated by vagal sensory neurons expressing substance P, calcitonin gene-related peptide (CGRP), and 5-HT3 receptors^7^. However, there is limited information available about the molecular identity of vagal sensory neurons that innervate the liver and their role in hepatic metabolism. It appears that only a limited number of vagal sensory neurons in the left nodose ganglion project to the liver in rats^11^. The peripheral nerve terminals of the vagal sensory neurons innervating the liver are primarily present in the bile ducts and portal veins, whereas no afferent terminals are observed in the hepatic parenchyma of rats^12^.

The liver is the largest metabolic organ responsible for the body’s energy needs and surfeits. As the functions of organs, including the brain, depend largely on the fuel produced by the liver, dysregulation of hepatic metabolism would significantly affect their functions. Although the brain is the most energy-demanding organ in the body, it remains relatively unknown whether hepatic metabolism can influence brain function. In fact, individuals with metabolic dysfunction-associated fatty liver disease (MAFLD) have a higher risk of psychiatric disorders including anxiety, depression, bipolar disorder, schizophrenia, and dementia^13–15^, suggesting that impaired hepatic lipid metabolism may be closely associated with mental illness. As liver-derived interoceptive signaling molecules can reach higher brain centers via hepatic vagal sensory neurons and humoral pathways, it is reasonable to assume that disrupted interoceptive signaling caused by MAFLD may result in psychiatric illness.

In this study, we sought to determine the molecular identity of vagal sensory neurons innervating the liver and their role in controlling hepatic steatosis and anxiety-like behavior. We found that a small subset of polymodal vagal sensory neurons projected to the liver. These sensory neurons played an indispensable role in the development of hepatic steatosis in mice fed a high-fat diet (HFD) as the loss of the liver-brain axis prevented diet-induced obesity (DIO) and hepatic steatosis in both female and male mice fed HFD. In addition, mice lacking liver-projecting vagal sensory neurons exhibited a significant reduction in anxiety-like behavior, suggesting an important role of liver-innervating vagal sensory neurons in developing psychiatric and metabolic disorders associated with obesity and diabetes.

## Results

### Molecular identification of vagal sensory neurons innervating the liver

To determine the degree to which vagal sensory neurons innervate the liver, we expressed the Cre recombinase in the vagus nerve ganglia by administering retrograde adeno-associated viruses (AAVrg) encoding the Cre gene (AAVrg-Cre) to the liver of Rosa26-eGFP^f^ mice (Fig. 1a). Under these conditions, AAVrg-Cre viruses were taken up by the peripheral vagal nerve terminals and transported toward their respective cell bodies in the nodose ganglia. GFP immunostaining revealed the presence of GFP-positive cells in the vagus nerve ganglion (Fig. 1b). Contrary to previous studies showing that vagal sensory neurons innervating the liver are primarily located in the left nodose ganglion in rats^111,16^, the number of GFP-positive neurons was similar in both vagus nerve ganglia (Fig. 1c).

**Figure 1.**
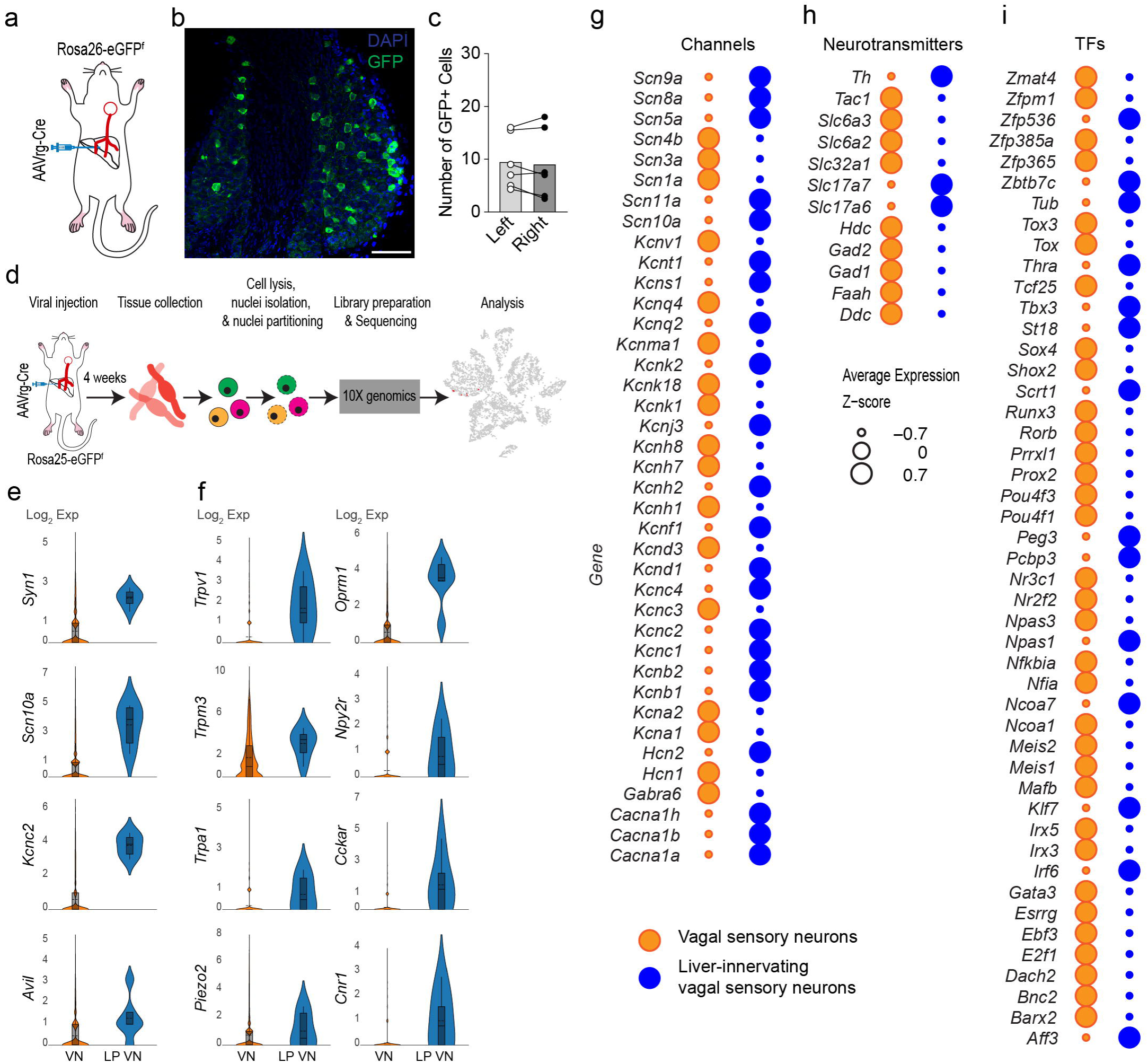
Molecular identification of vagal sensory neurons innervating the liver. **a.** Schematic illustration of our experimental configurations. AAVrg-Cre viruses were injected into the medial and left lobes of the livers of Rosa26-eGFP^f^ mice. **b.** Image from confocal microscopy showing GFP-positive cells in the right nodose ganglion of Rosa26-eGFP^f^ mice injected with AAVrg-Cre into the liver. Scale bar, 100 μm **c.** Graph showing the number of GFP-positive cells per section in the left and right nodose ganglia. **d.** Schematic illustration depicting our experimental process. **e** and **f.** Violin plots showing enriched genes in liver-projecting vagal sensory neurons (LP VN) compared to other vagal sensory neurons (VN) (neuronal marker genes (e), membrane receptors (f). **g**, **h**, and **i**. Bubble heatmaps showing differentially expressed genes in liver-projecting vagal sensory neurons compared to other vagal sensory neurons (voltage-gated channels (g), neurotransmitters (h), and TFs (i).

We then sought to determine the molecular identities of liver-innervating vagal sensory neurons by performing snRNA-Seq of vagal sensory neurons in Rosa26-eGFP^f^ mice injected with AAVrg-Cre into the liver (Fig. 1d). We recently identified approximately 9,000 cells from 30 nodose ganglia collected from 8 males and 7 females^17^. Among them, approximately 6,000 cells had neuronal marker genes, including *Map2*, *Syn1, Syn2*, and *Slc17a6*. However, we identified only 6 genetically tagged vagal sensory neurons (Supplementary Table 1) because of the technical limitations associated with a limited number of liver-innervating vagal sensory neurons. We grouped the six vagal sensory neurons innervating the liver and compared them with other vagal sensory neurons. They expressed neuronal marker genes such as synapsin 1 (*Syn1*), voltage-gated sodium channel α subunit 10 (*Scn10a*), voltage-gated potassium channel subfamily C member 2 (*Kcnc2*), and the sensory neuron-specific actin-binding protein advillin (*Avil)*^18^. They also expressed several ion channel subtypes including chemosensitive transient receptor potential cation channel subfamily V member 1 (*Trpv1*), temperature-sensitive TRP subfamily M member 3 (*Trpm3*) and subfamily A member 1 (*Trpa1*), and mechanosensitive piezo-type ion channel component 2 (*Piezo2*) (Fig. 1f), indicating that they are polymodal sensory neurons. These neurons also expressed the opioid receptor μ-1 (*Oprm1*), the neuropeptide Y receptor type 2 (*Npy2R*), the cholecystokinin A receptor (*Cckar*), and the cannabinoid receptor type 1 (*Cnr1*).

Intriguingly, vagal sensory neurons innervating the liver exhibited high expression levels of *Hcn2*, *Kcns1*, *Scn9a*, *Scn10a*, and *Scn11a* when compared to other vagal sensory neurons (Fig. 1g). Notably, these voltage-gated channels are primarily observed in polymodal nociceptors located in the dorsal root ganglion^19–21^. The present liver-innervating vagal sensory neurons were predominantly glutamatergic, expressing the *Slc17a6* gene (Fig. 1h). We also noted that liver-innervating sensory neurons highly expressed genes for transcription factors (TFs) associated with energy metabolism and behaviors, including *Ncoa7*, *Thra,* and *Tbx3* genes when compared to other vagal sensory neurons (Fig. 1i). It has been well described that thyroid hormones regulate feeding, thermogenesis, systemic glucose and lipids levels^22^ and that loss of the hypothalamic *Tbx3* gene caused an increase in food intake and body weight^23^. Hence, our initial molecular analysis demonstrates that vagal sensory neurons innervating the liver exhibit polymodal molecular genetic identities characteristic of nociceptive sensory neurons.

### The liver is innervated by a subset of Avil-positive vagal sensory neurons

Based on our snRNA-Seq results and a previous study indicating that chemosensitive vagal sensory neurons express the *Avil* gene^2^, we used the Avil^CreERT2^ strain to identify vagal sensory neurons that innervate the liver. Additionally, we employed AAVrg-FLEX-jGCamp7s, which expresses Cre-dependent jGCamp7s, due to our observation that GFP expression with this vector was more robust compared to other AAV vectors encoding eGFP and tdTomato.

Administering AAVrg-FLEX-jGCamp7s to the medial and left lobes of the livers of Avil^CreERT2^ mice resulted in the selective labeling of liver-innervating vagal sensory neurons (Fig. 2a, b, and c). Both left and right nodose ganglia exhibited a small number of GFP-positive cells. Approximately 10% of Avil-positive vagal sensory neurons were positive for GFP. We subsequently examined whether these Avil-positive vagal sensory neurons send axonal projections to the liver by employing a retrograde AAV that encodes a Cre-dependent placental alkaline phosphatase (PLAP), which is expressed on the plasma membrane (AAVrg-FLEX-PLAP) (Fig. 2d). Injection of this AAVrg-FLEX-PLAP into the liver of Avil^CreERT2^ mice resulted in the presence of PLAP-positive nerves in the parenchyma of the liver, particularly in the periportal and midlobular areas (Fig 2e).

**Figure 2.**
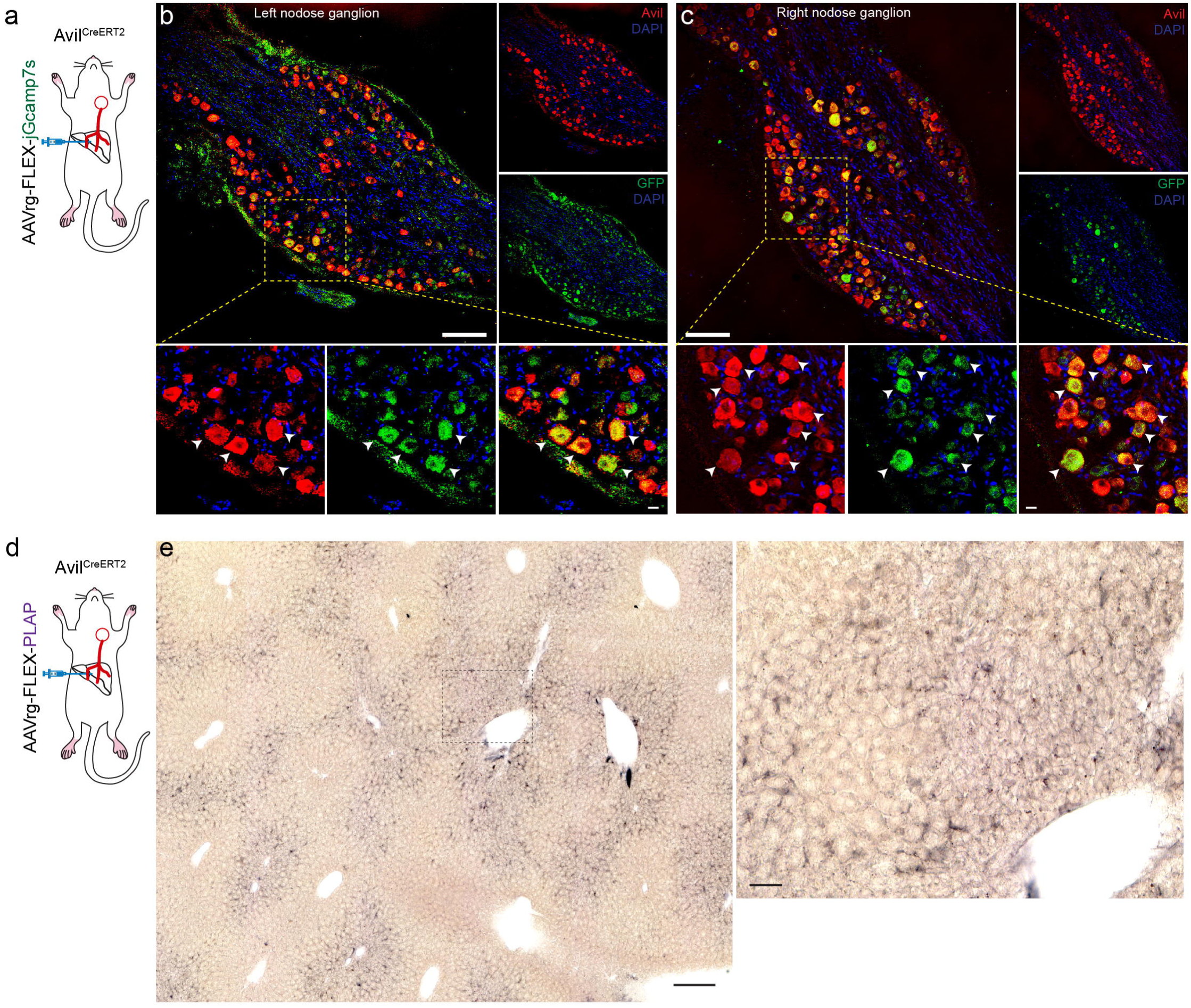
Immunohistochemical identification of vagal sensory neurons innervating the liver. **a.** AAVrg-FLEX-jGcamp7s viruses were injected into the livers of Avil^CreERT2^ mice. **b and c**. Images showing co-expression of GFP and Avil in a subset of neurons in the left and right nodose ganglia of Avil^CreERT2^ mice injected with AAVrg-FLEX-jGcamp7s into the liver. Upper panel: scale bar, 100 μm. Bottom panel: higher magnification view of the yellow square area. Arrowheads represent the vagal sensory neurons that innervate the liver. Scale bar, 10 μm **d** and **e.** Schematic illustration of our experimental conditions (d). Images showing PLAP-positive nerve fibers in the liver parenchyma. Right panel: Higher-magnification view of the black square box in the left panel. Scale bars, 200 μm (left), 50 μm (right)

To further investigate their peripheral nerve terminals of these liver-innervating vagal sensory neurons, we crossbred the Avil^CreERT2^ strain with floxed-Stop channelrhodopsin (ChR2)- tdTomato mice (Avil^CreERT2:ChR2-tdTomato^), as the ChR2-tdTomato fusion protein displayed strong expression in nerve terminals (Fig. 3a). Immunostaining revealed numerous tdTomato-positive nerve fibers in the periportal area (Fig. 3b), indicating that they were the nerve fibers originated in Avil-positive sensory neurons. Next, we administered anterograde AAV1-FLEX-GFPsm-Myc directly into the left nodose ganglion of Avil^CreERT2^ mice (Fig. 3c and d) to label peripheral hepatic nerve terminals via anterograde viral transport. We found the anterogradely-labeled GFP- and Myc-positive nerve terminals primarily around the periportal, but not pericentral, areas (Fig. 3d and Supplementary Fig. 1a and b). We further examined their peripheral nerve terminals by injecting retrograde AAV (AAVrg)-FLEX-jGCamp7s into the livers of Avil^CreERT2^ mice (Fig. 3e and Supplementary Fig. 1c). Avil-positive nerve terminals were co-labeled with an anti-GFP antibody in the periportal area (Fig. 3f), whereas there were no Avil-positive nerves in the pericentral zone (Supplementary Fig. 1d). Additionally, co-staining with an antibody against the presynaptic marker synapsin 1 (Syn1) revealed that GFP-positive nerve terminals were positive for Syn1, indicating that these are nerve terminals (Fig. 3g). Taken together, our tracing results strongly support the interpretation that the liver receives synaptic input from Avil-positive vagal sensory neurons.

**Figure 3.**
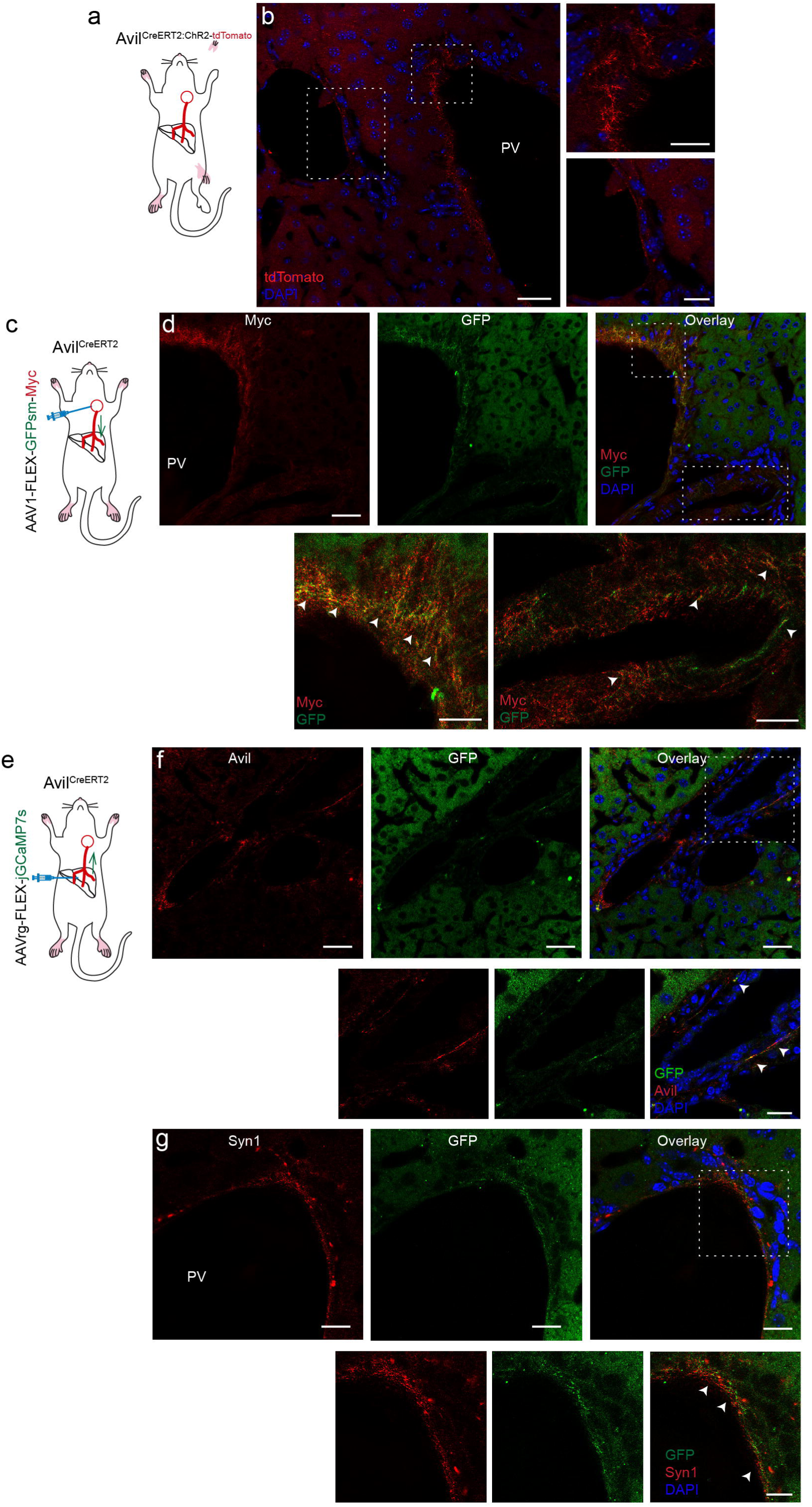
Avil-positive vagal sensory neurons send axonal projections to the periportal area. **a** and **b**. Schematic illustration of the experimental conditions. We used Avil^CreERT2;ChR2-tdTomato^ mice. Images showing expression of tdTomato at nerve terminals in the periportal area (b). Left panel, scale bar, 20 μm. Right panel: higher magnification view of the white square area, scale bar, 10 μm **c** and **d.** Anterograde monosynaptic AAV-FLEX-GFPsm-myc viruses were injected into the left nodose ganglion of of Avil^CreERT2^ mice. Most nerve terminals were positive for GFP and Myc, indicating that they were the nerve terminals of Avil-positive vagal sensory neurons innervating the liver. Scale bar, 20 μm. Bottom panel: higher magnification view of the white square area. Arrowheads represent co-expression of Myc and GFP. Bottom panel: scale bar, 10 μm **e.** AAVrg-FLEX-jGcamp7s viruses were injected into the livers of Avil^CreERT2^ mice. **f** and **g**. Images showing that GFP-positive nerve terminals were also stained with Avil (red, f) and the presynaptic marker Syn1 (red, g). Upper panel: scale bar, 20 μm, Bottom panel: scale bar, 10 μm

We then examined the medullary CNS projection sites of the hepatic vagal sensory neurons using brainstem sections from Avil^CreERT2^ mice injected with AAVrg-FLEX-GFP into the liver (Supplementary Fig. 2a). Following the tissue-clearing process, we conducted a 3D imaging analysis of the tissues, which revealed GFP-positive puncta and nerve terminals in the NTS and the area postrema (AP) (Supplementary Fig. 2b and Supplementary Movie 1). Additionally, scattered GFP-positive puncta were observed in the dorsal motor nucleus of the vagus (DMV). (Supplementary Fig. 2b and Supplementary Movie 1).

### Loss of liver-innervating vagal sensory neurons causes weight loss and increases energy expenditure

The hepatic portal vein carries nutrients and hormones from the gastrointestinal tract^24^. As vagal sensory neurons sent projections to the periportal regions of the liver and expressed lipid-sensing TRP receptors (Fig. 1f), it is possible that vagal sensory neurons innervating the liver may detect changes in nutrients^25^. We thus investigated the impact of the loss of the liver-brain axis on energy balance under obesogenic conditions. To ablate liver-innervating vagal sensory neurons, we expressed genetically engineered taCaspase-3 (taCasp3)^26^ in the vagal sensory neurons innervating the liver by injecting AAVrg-FLEX-taCasp3-TEVp into the livers of Avil^CreERT2^ mice (Fig. 4a). To determine whether the retrograde expression of taCasp3-TEVp ablates liver-projecting vagal sensory neurons, we administered the retrograde tracer Fluoro-Gold into livers of Avil^CreERT2^ mice injected with AAVrg-FLEX-taCasp3-TEVp and examined the number of Fluoro-Gold-positive vagal sensory neurons (Supplementary Fig. 3a). There was a significant reduction of liver-projecting vagal sensory neurons (Supplementary Fig. 3b, c, and d). Moreover, unlike the Avil^CreERT2^ mice injected with AAVrg-FLEX-jGCamp7s, we did not observe any GFP-positive nerve terminals in the periportal areas of AAVrg-FLEX-taCasp3-TEVp injected mice (Supplementary Fig. 3e, f, and g), which provides further evidence of the loss of liver-projecting vagal sensory innervation of the liver.

**Figure 4.**
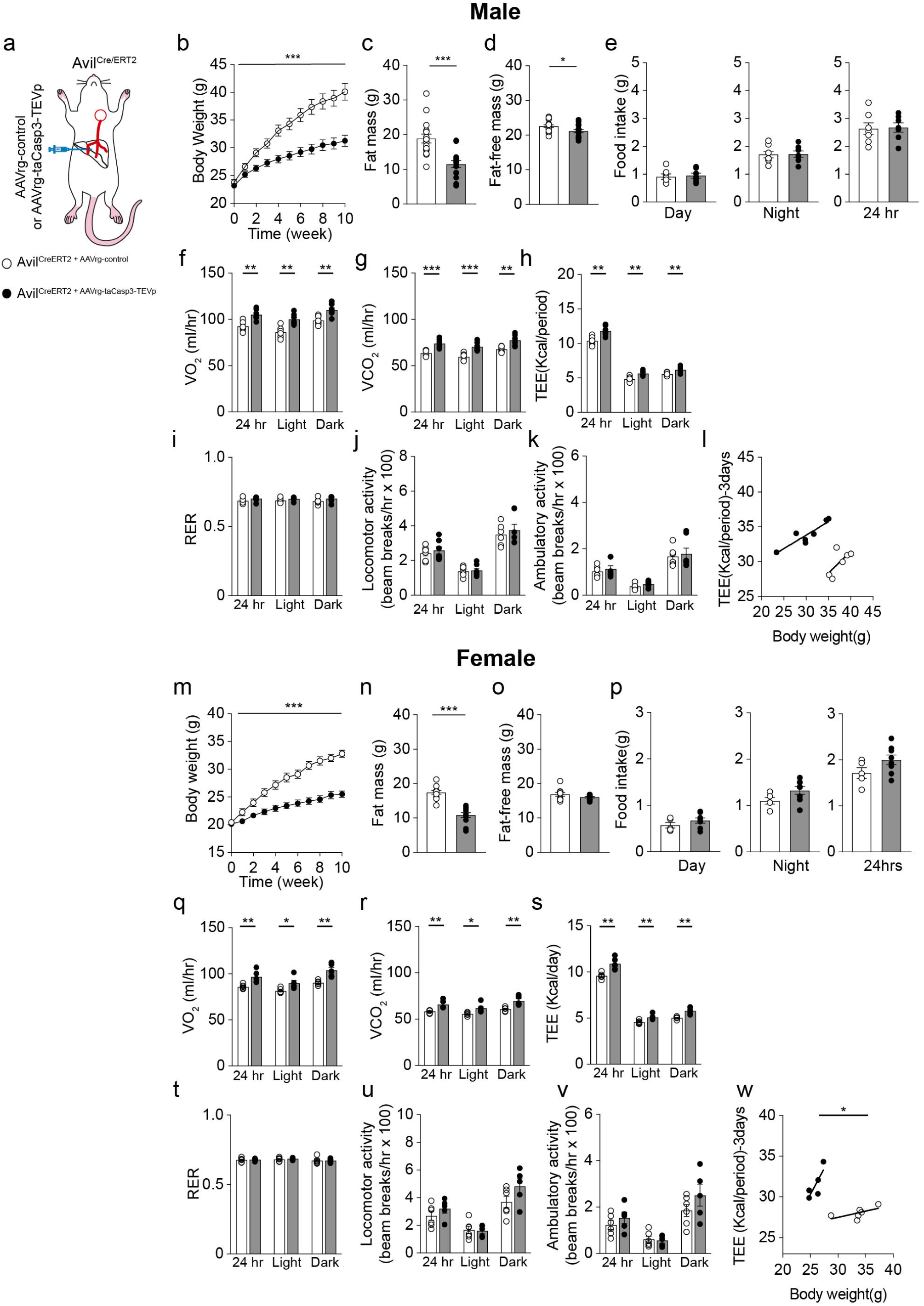
Loss of the liver-brain axis reduces body weight and increases energy expenditure in mice fed HFD. **a.** Schematic representation of the experimental configurations. AAVrg-FLEX-eGFP and AAVrg-FLEX-taCasp3-TEVp viral vectors were injected into the livers of Avil^CreERT2^ mice. **b.** Graph showing the changes in body weight of the control (n= 16 mice) and experimental groups (n= 15 mice) (two-way ANOVA, ***p<0.001) **c and d**. Graphs showing the differences in fat mass and fat-free mass between the two groups (control, n= 16 mice; experimental, n=15 mice; unpaired *t*-test, ***p<0.001, *p<0.05). **e**. Graphs showing no difference in food intake between the two groups (control, n= 6 mice; experimental mice, n=7 mice). **f** and **g.** Graphs showing an increase in O_2_ consumption and CO_2_ production in the experimental group compared to the control group (control, n= 6 mice; experimental mice, n=7 mice; unpaired *t*-test, **p<0.01, ***p<0.001). **h**. Graph showing that the experimental mice exhibited a significant increase in TEE compared with the control group (control, n= 6 mice; experimental mice, n=7 mice; unpaired *t*-test, **p<0.01). **i, j, and k**. Graphs showing the RER and locomotor activity (control, n= 6 mice; experimental mice, n=7 mice). **l**. ANCOVA analysis showing no difference between the two groups (control, n= 6 mice; experimental mice, n=7 mice). **m**. Graph showing the changes in body weight of the control and experimental groups (control, n= 15 mice; experimental mice, n= 21 mice; Two-way ANOVA, ***p<0.001). **n and o**. Graphs show that the experimental mice exhibited lower fat mass than the controls (control, n= 15 mice; experimental mice, n= 21 mice; unpaired *t*-test, ***p<0.001). **p**. Graphs showing no difference in food intake between the two groups (control, n= 5 mice; experimental mice, n= 8 mice). **q** and **r.** Summary graph showing an increase in O_2_ consumption and CO_2_ production in the experimental group compared to the control group (control, n= 6 mice; experimental mice, n= 5 mice; unpaired *t*-test, *p<0.05, **p<0.01). **s**. Graph showing that the experimental mice displayed a greater TEE than the control group (control, n= 6 mice; experimental mice, n= 5 mice; unpaired *t*-test, **p<0.01). **t, u, and v**. Graphs showing no differences in RER or locomotor activity between the groups (control, n= 6 mice; experimental mice, n= 5 mice). **w**. ANCOVA analysis showing a significant difference in the slope between the groups (control, n= 6 mice; experimental mice, n= 5 mice, *p<0.05).

Under these experimental conditions, measurements of body weight showed a significant difference in body weight between the groups. The experimental male mice gained significantly less weight than control mice (Fig. 4b). This finding aligns with a recent study demonstrating that the elimination of vagal sensory neurons innervating the liver prevents DIO ^27^. This was accompanied by a significant decrease in both fat mass and fat-free mass in the experimental mice compared with the controls (Fig. 4c and d). We then examined whether changes in food intake and/or energy expenditure in male mice fed HFD result in decreased body weight. Both groups consumed similar amounts of food during the light and dark phases (Fig. 4e). We thus investigated if there was a difference in the energy expenditure between the groups. The experimental mice consumed more O_2_ and produced more CO_2_ than the control mice (Fig. 4f and g), resulting in a significant increase in total energy expenditure (TEE) over a 24-hour period (Fig. 4h). The respiratory energy ratio (RER) and locomotor activity did not differ between the groups (Fig. 4i, j, and k). Analysis of covariance (ANCOVA) using body weight as a covariate showed no differences in the slopes (Fig. 4l), suggesting that body fat did not have a substantial impact on TEE.

Ablating liver-innervating vagal sensory neurons caused a reduction in body weight in female mice as well (Fig. 4m). This reduction was primarily associated with decreased fat mass but not fat-free mass (Fig. 4n and o). Like males, female mice lacking hepatic vagal sensory neurons displayed no difference in food intake compared to controls (Fig. 4p). Indirect calorimetry analysis revealed that VO_2_, VCO_2_, and TEE in experimental mice were significantly higher than those in control mice (Fig. 4q, r, and s). There were no differences in the RER and locomotor activity between the two groups (Fig. 4t, u, and v). Intriguingly, ANCOVA analysis revealed that body weight significantly influenced TEE (Fig. 4w), suggesting that fat tissue may significantly impact energy expenditure in females.

### Loss of liver-innervating vagal sensory neurons improves glucose tolerance in mice fed HFD

We next examined the impact of the loss of the liver-brain axis on glucose homeostasis during HFD feeding. Measuring the glucose levels following 10 weeks of HFD feeding revealed that both non-fasting and fasting blood glucose levels were significantly lower in the experimental male mice than in the control male mice (Supplementary Fig. 4a and b). Intraperitoneal glucose tolerance tests (i.p.GTT) showed that the blood glucose levels of the experimental mice in response to a bolus of glucose (i.p., 2 g/kg) were significantly lower than those of the control mice (Supplementary Fig. 4c and d). Insulin tolerance tests (ITT 1 U /kg, i.p.) revealed improved tolerance in experimental males compared to control mice (Supplementary Fig. 4e). This improved insulin tolerance in the experimental group was associated with decreased basal and fasting insulin levels relative to those in the control group (Supplementary Fig. 4f and g).

Likewise, female mice lacking hepatic vagal sensory neurons exhibited lower basal and fasting blood glucose levels than the control females (Supplementary Fig. 4h and i). Female mice in the experimental group displayed improved glucose tolerance during the GTT compared to the control group (Supplementary Fig. 4j and k). However, there was no noticeable difference in insulin sensitivity between the two groups during the ITT (Supplementary Fig. 4l). The basal insulin levels were significantly lower in the experimental group compared to the control group, while the fasting insulin levels were similar (Supplementary Fig. 4m and n).

### Loss of liver-innervating vagal sensory neurons prevents hepatic steatosis in HFD-fed mice

We then investigated whether the loss of the liver-brain axis affects hepatic lipid metabolism. The livers of the control group turned yellowish, while the livers of the experimental mice remained red (Fig. 5a). The liver weight was significantly lower in the experimental mice than in the control mice (Fig. 5b). Hematoxylin and Eosin (H&E) staining revealed clear differences between the groups (Fig. 5c). Specifically, mice displayed numerous large and small lipid droplets, primarily in the pericentral regions, compared to experimental mice. Oil Red O staining further confirmed hepatic fat accumulation in control mice (Fig. 5d).

**Figure 5.**
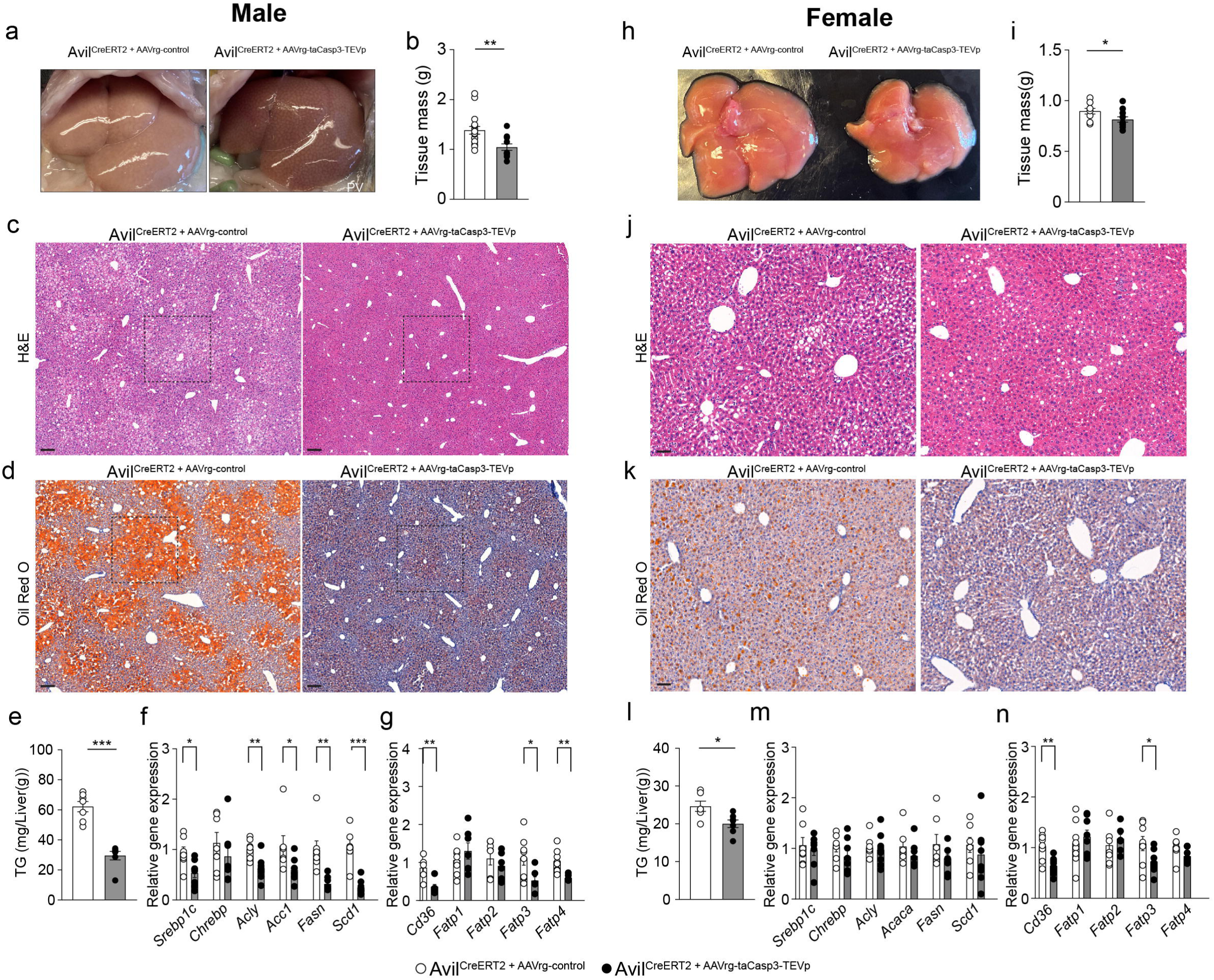
Loss of the liver-brain axis prevents hepatic steatosis in mice fed HFD. **a.** Representative images of liver tissues from the control and experimental mice. **b.** Graphs showing liver mass between the groups (n= 14 vs. 18 mice, unpaired *t*-test, **p<0.01) **c and d. .** H&E (c) and Oil Red O (d) staining of liver tissues from the control and experimental mice. Nuclear staining with hematoxylin (blue), Scale bar, 100 μm. **e.** Graph showing hepatic TG levels between the groups (n= 7 mice vs. 7 mice, unpaired *t*-test, ***p<0.001). **f.** Summary graphs showing changes in hepatic mRNA expression of enzymes involved in lipid metabolism between the groups (control, n= 7; experimental, n= 7, unpaired t-test, *p<0.05, **p<0.01, ***p<0.001). **g.** Summary graphs showing changes in hepatic mRNA expression of fatty acid uptake transports between the groups (control, n= 6-8 mice; experimental, n= 7 mice, unpaired t-test, *p<0.05, **p<0.01). **h.** Images of liver tissues from the control and experimental mice. **i.** Graphs showing that the liver mass of the experimental mice (n=9) was lower when compared to that of the control group (n= 11 mice, unpaired *t*-test, *p<0.05). **j and k.** H&E (j) and Oil Red O (k) staining of liver tissues from the control and experimental mice. Scale bar, 50 μm. **l.** Graph showing hepatic TG levels between the groups (n= 6 mice vs. 7 mice, unpaired *t*-test, *p<0.05). **m.** Summary graph showing alterations in hepatic lipogenic gene expression levels (control, n= 7; experimental mice, n= 8). **n.** Summary graph showing changes in hepatic fatty acid uptake transport expression levels (control, n= 8; experimental mice, n= 8, unpaired t-test, *p<0.05, **p<0.01).

Given the lack of lipid droplets in the livers of experimental male mice, we measured hepatic triglyceride (TG) levels. The experimental mice displayed a substantial decrease in hepatic TG levels compared with the control mice (Fig. 5e). RT-qPCR analysis of liver tissues from the experimental mice revealed a significant decrease in the expression of the lipogenic transcription factor *Srebp-1c*, in comparison to the controls (Fig. 5f). This was accompanied by downregulation of its target genes, including *Acaca*, *Fasn*, and *Scd1* (Fig. 5f). In addition, the impairment of the liver-brain axis downregulated the fatty acid translocase *Cd36* and the fatty acid transmembrane transporters *Fatp3* and *Fatp4* gene expression (Fig. 5g). These results indicate that the loss of the liver-brain axis may lead to a reduction in *de novo* lipogenesis (DNL) and fatty acid uptake, thereby preventing the onset of hepatic steatosis in male mice fed HFD.

In female mice, there was no visible difference in liver color (Fig. 5h). The liver weight was slightly but significantly reduced in the experimental group compared to the control group (Fig. 5i). H & E and Oil red O staining revealed that the control group exhibited small lipid droplets in the liver parenchyma, whereas the experimental group displayed fewer such droplets (Fig. 5j and k). The experimental mice demonstrated a decrease in hepatic TG compared to the control mice (Fig. 5l). In contrast to the experimental male mice, the experimental female mice showed no significant difference in the expression of the lipogenic genes (Fig. 5m). Nonetheless, female mice lacking liver-innervating vagal sensory neurons exhibited decreased *Cd36* and *Fatp3* expression levels (Fig. 5n). Taken together, these novel findings support the interpretation that liver-innervating vagal sensory neurons are crucial for the emergence of hepatic steatosis in mice during the development of obesity.

The male control mice displayed dense lipid accumulation in the pericentral areas compared to the experimental mice. We conducted further analysis to determine if there are distinct gene expression patterns throughout the hepatic zonation in both groups by performing spatial transcriptomics (Fig. 6). Pericentral, midlobular, and periportal clusters were identified based on the known zonation markers (Fig. 6a). We found that approximately 500 genes in the PC and PP areas were significantly different between the groups (Fig. 6b and Supplementary Table 2 and 3) and that the midlobular areas exhibited about 300 differentially expressed genes (DEGs) (Fig. 6b and Supplementary Table 4). We conducted a selective gene ontology (GO) analysis of the DEGs in each zonation using Enrichr-KG^28^, which identified the top 5 enriched GO terms in biological process (Fig. 6c). These included “cotranslational protein targeting to membrane” (GO:0006613), “SRP-dependent cotranslational protein targeting to membrane” (GO:0006614), “protein targeting to ER” (GO:0045047), “cytoplasmic translation” (GO:0002181), and “cellular protein metabolic process” (GO:0044267). According to mammalian phenotype ontology analysis^29^, each zonation shared 4 terms, including “abnormal lipid homeostasis” (MP:0002118), “abnormal liver physiology” (MP:0000609), and “hepatic steatosis” (MP:0002628). Notably, each zonation had a unique GO term, such as “increased liver weight” (MP:0002981) in the PP group, “abnormal hepatocyte morphology” (MP:0000607) in the Mid group, and “abnormal circulating amino acid level” (MP:0005311) in the PC group. Furthermore, the Jesensen diseases ontology analysis revealed a term for “fatty liver disease” (Fig. 6c). These results further support the interpretation that liver-projecting vagal sensory neurons play an important role on developing hepatic steatosis in mice fed HFD.

**Figure 6.**
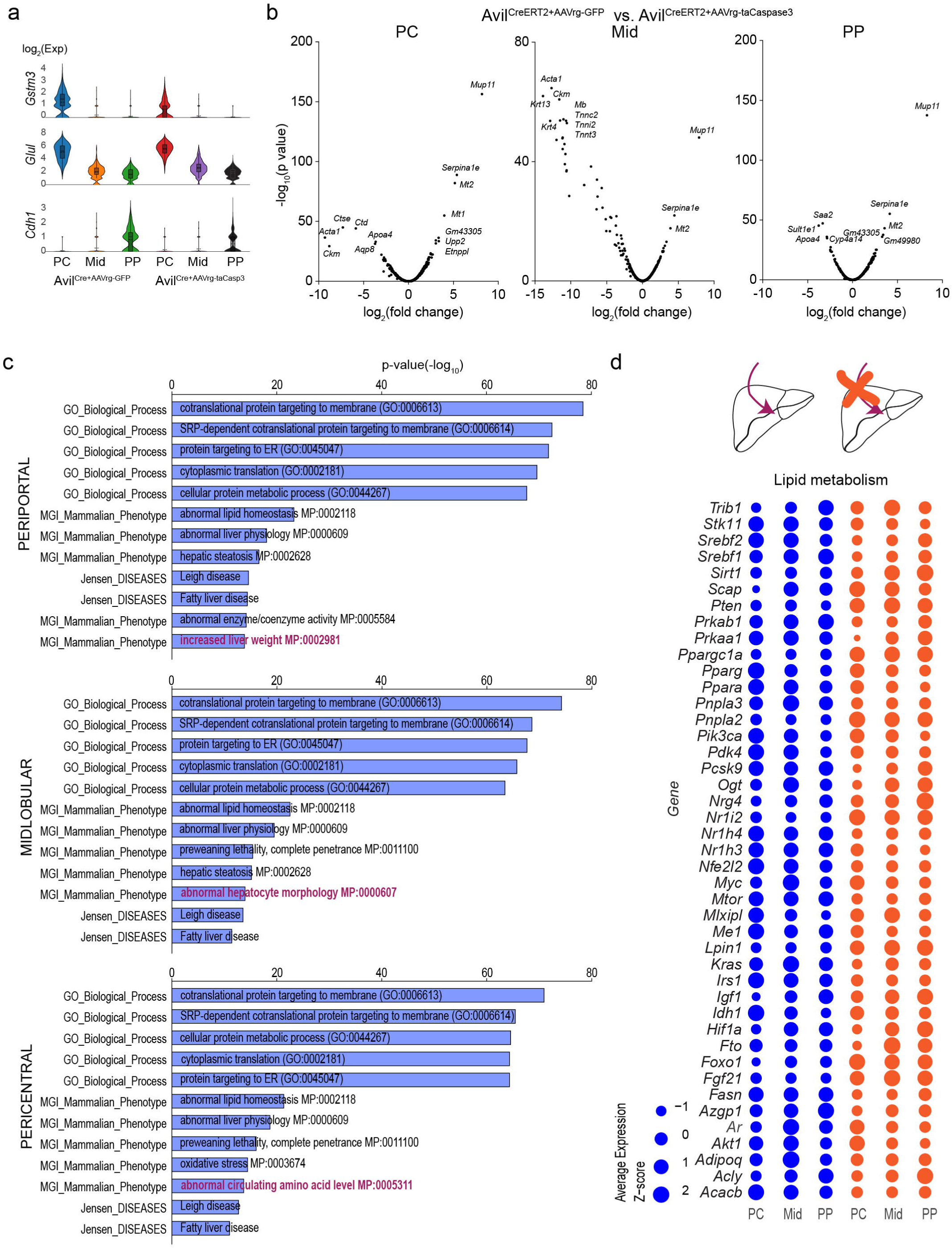
Differential gene expression in the livers of mice with and without liver-innervating vagal sensory neurons. **a.** Violin plots depicting the differential expression of the liver zonation marker genes in the pericentral, midlobular, and periportal areas between the groups. **b.** Volcano plots illustrate the differences in gene expression across the zonation of the livers between the groups. **c.** Plots displaying the top GO terms in biological process, mammalian phenotype, and Jensen diseases across the zonation of the livers between the groups. **d.** Bubble heatmap showing differentially expressed genes associated to lipid metabolism between mice with and without liver-innervating vagal sensory neurons.

Our GO analysis highlighted significant changes in genes linked to hepatic lipid metabolism. We thus sought to identify the genes affected by the deletion of hepatic vagal sensory neurons. We found that the loss of liver-projecting vagal sensory neurons resulted in the opposite regulation of two transcription factors, specifically *Irf6* and *Tbx3* (Supplementary Fig. 5a and b). It has been described that HFD feeding downregulated *Irf6* expression in hepatocytes and that overexpression of *Irf6* in hepatocytes reduced hepatic steatosis^30^. In contrast, deletion of *Tbx3* improved hepatic steatosis in mice fed a high-fat diet^31^. Mice in the control group demonstrated increased expression of *Acacb*, *Fasn*, *Me1*, *Pcsk9*, *Srebf1*, and *Srebf2* genes (Fig. 6d). Conversely, mice lacking hepatic vagal sensory input showed enhanced expression of *Lpin1*, *Pnpla2*, and *Ppargc1a* genes (Fig. 6d). Additionally, mice lacking hepatic sensory neurons exhibited reduced expression of the glucokinase (GCK) gene (Supplementary Fig. 5c). This gene is responsible for the phosphorylation of glucose to produce glucose 6-phosphate, which subsequently triggers hepatic lipogenesis^32^. In addition, the loss of hepatic sensory input elevated *Fgf21*, *Igfbp1*, *Igfbp2*, and *Igfbp4* genes, which downregulated lipogenesis^33,34^ (Supplementary Fig. 5d). Taken together, these results further support the interpretation that mice lacking hepatic sensory input exhibited reduced fatty acid synthesis and cholesterol, along with enhanced β-oxidation and lipolysis.

### Loss of the liver-brain axis improves anxiety-like behavior in male mice fed HFD

Individuals with MAFLD have been found to have an increased risk of developing psychiatric disorders, including anxiety and depression^13–15^. To investigate if the vagal sensory pathway that innervates the liver also affects animal behavior, particularly anxiety-related behavior, we conducted a series of experiments. In the open field exploration test, which is an initial screening for anxiety-like behavior^35^, experimental male mice displayed increased time spent in the inner area and decreased time spent in the outer area (Fig. 7a and b). We then performed an elevated plus maze (EPM) test that examines the natural tendency of mice to explore novel environments by giving the choice of spending time in open, unprotected maze arms or enclosed, protected arms^36^ (Fig. 7c). The control mice spent more time in the closed arm (Fig. 7c) and made fewer entries into the open arms than the experimental group. The light-dark (LD) box test that assesses a change in willingness to explore the illuminated, unprotected area revealed that the experimental mice exhibited increased time spent in the light area compared to the control group (Fig. 7d). While the female experimental mice displayed increased locomotor activity, increased time spent and zone alteration (Fig. 7g), they did not show any significant differences in the elevated plus maze (EPM) or LD box test (Fig. 7h an i). Additionally, we evaluated depression-like behavior using the forced swim test and tail suspension test. Our findings revealed no significant differences in these assessments for both male and female mice (Fig. 7e, f, j, and k). The majority of these results support the interpretation that mice lacking liver-projecting vagal sensory neurons exhibited significant improvements in anxiety-like behavior in male mice.

**Figure 7.**
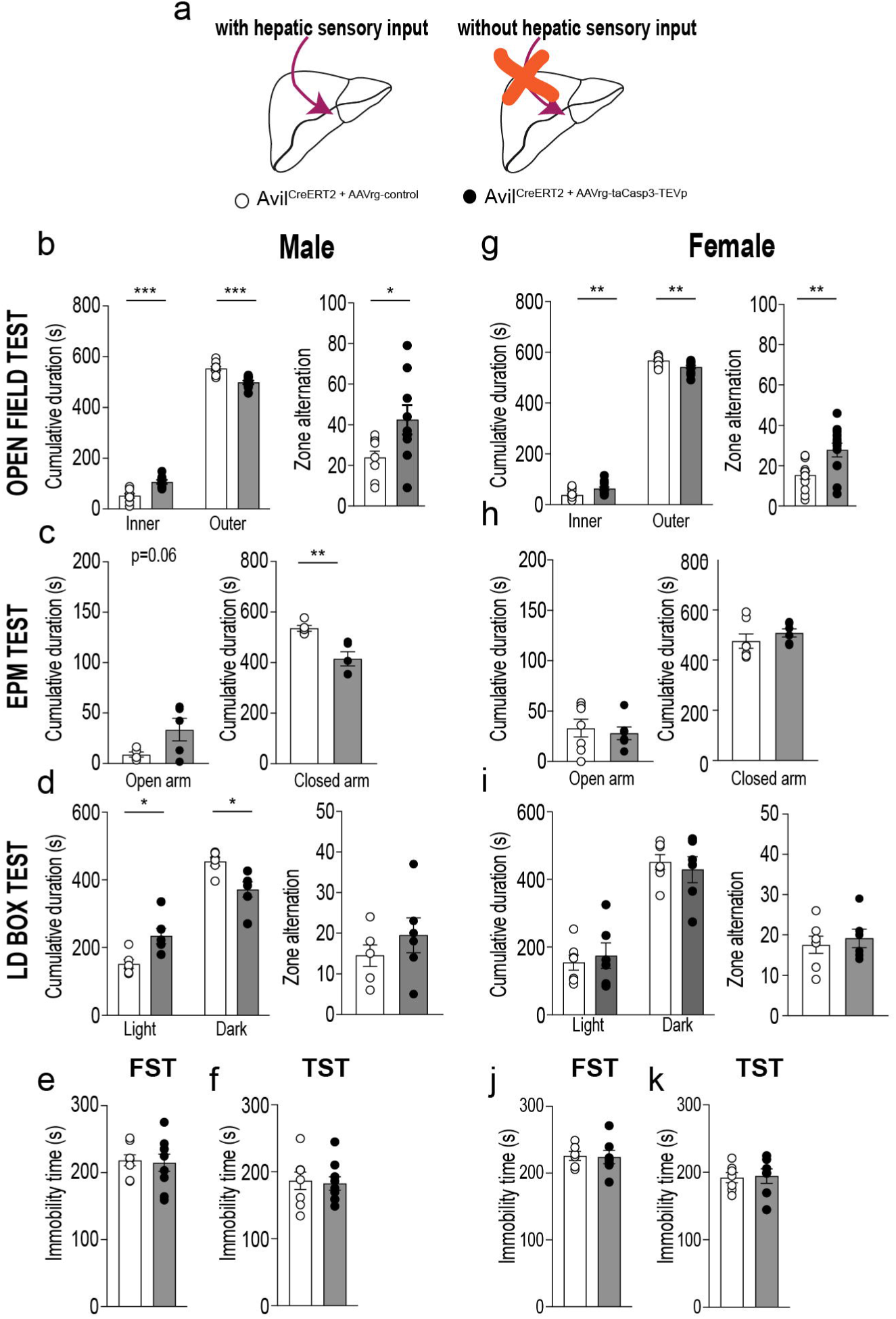
Deleting liver-innervating vagal sensory neurons improves anxiety-like behavior. **a.** Schematic representation of the experimental configurations. **b.** Graphs showing the time spent in the inner and outer areas and zone alteration in the open filed test between the male groups (n= 9 mice vs. 9 mice, two-tailed t-test, *p<0.05, ***p<0.001) **c.** Graphs showing the cumulative duration spent in the open and closed arms in the EPM test between the male groups (n= 5 mice vs. 5 mice, two-tailed t-test, *p<0.05, **p<0.01) **d.** Graphs showing the time spent in the light and dark zones and the number of zone transition for control (n= 6 mice) and experimental (n= 6 mice) male mice during the light-dark box assessment. Two-tailed t-test, *p<0.05 **e** and **f**. Graphs showing that the FST and TST evaluations revealed no significant differences in immobility time between the groups (n= 8 mice vs. 9 mice). g. Graphs showing the time spent in the inner and outer areas and zone alteration in the open filed test between the female groups (n= 13 mice vs. 12 mice, two-tailed t-test, **p<0.01). h, I, j, and k. Graphs showing no significant differences between females with and without hepatic vagal sensory input across several behavioral assessments. These tests included the EPM test (h, n=7 mice vs. 6 mice), LD box test (I, n=7 mice vs. 6 mice), FST (j, n=6 mice vs. 7 mice), and TST (k, n=7 mice vs.7 mice).

## Discussion

This study provides molecular, anatomical, functional, and behavioral evidence for the pivotal role of the liver-brain axis in controlling energy balance, glucose homeostasis, hepatic steatosis, and anxiety-like behavior in mice fed obesogenic HFD. Our single-nucleus RNA-seq analysis revealed that liver-projecting vagal sensory neurons were polymodal C-fiber sensory neurons as previous described^2^ (i.e., NG12 to NG18). In fact, polymodal C-fiber vagal sensory neurons expressed *Scn10a*, *Trpv1, Trpa1*, *Npy2r*, and *Cckar*^2^ which were also expressed in liver-projecting vagal sensory neurons. Liver-innervating vagal sensory neurons were found mainly in the periportal area surrounding the hepatic arteries, bile ducts, and portal veins and expressed mechano-sensing Piezo channels^37^, lipid-sensing TRPA1, TRPM3, and TRPV1 channels^38^, and gut-derived hormone CCK receptors. This multimodality of liver-innervating vagal sensory neurons would be particularly important because hepatic homeostasis highly relies on adequate perfusion and microcirculation via the dual hepatic arterial and portal venous blood supplies, as well as the fenestrated sinusoids. These anatomical and molecular features would enable them to accurately measure blood pressure, assess levels of various nutrients and peptides, and relay this information to the brain for processing and interpretation.

Intriguingly, our study revealed a significant expression of genes associated with lipid metabolism in liver-projecting vagal sensory neurons, such as *Acly*, *Fasn*, and *Scd2*. It has been shown that fatty acid metabolism in the brain, particularly the hypothalamus, exerts a significant influence on the regulation of food intake and energy metabolism^39–41^, suggesting that fatty acid metabolism and its associated metabolites in hypothalamic neurons may serve as indicators of nutrient status. Similarly, the increased lipogenic machinery in liver-innervating vagal sensory neurons may imply that they also possess the capability to control energy balance in response to changes in nutrient availability. Additionally, lipogenesis in neurons is critical for several aspects of neuronal function, including dendrite morphogenesis^42^, axon growth and regeneration^43^. The UNC5 family of netrin receptors (UNC5C and 5D) is involved in axon guidance and mediates axon repulsion of neuronal growth cones in the developing nervous system^44^. Of particular interest is that netrin-1 was found to stimulate the regeneration of the hepatic vagal nerve and the proliferation of hepatocytes in mice following hepatectomy^45^. Thus, our initial findings from single-nucleus RNA sequencing provides significant information regarding their potential implications for a range of critical physiological processes.

Prior retrograde labeling studies using horseradish peroxidase showed that a very small number of vagal sensory neurons in the left nodose projected to the liver in rats^11^. In contrast to this early finding, we found that both left and right nodose ganglia possessed similar numbers of liver-innervating vagal sensory neurons. A prior neuronal tracing study using a HSV viral tracer in rats showed that vagal sensory neurons projecting to the hepatic portal vein were also located in the left and right nodose ganglia of rats^46^. Furthermore, gut-innervating vagal sensory neurons were detected in both nodose ganglia as well^9^. It should be noted that the vagal sensory neurons innervating the liver traverse the common hepatic branch of the vagus nerve, which also includes those projecting to the portal vein and the gastroduodenal tract^47^. These prior studies support our results that liver-projecting vagal sensory neurons are located in the left and right nodose ganglia.

We report in this study that the periportal hepatic region received dense sensory innervation. As the portal vein delivers nutrients and hormones from the gut, periportal liver-projecting vagal sensory neurons appear to be ideally localized to transmit such gut-derived postabsorptive signals to the NTS. Both local interneurons and projection neurons in the NTS have been implicated in the control of various body functions. For instance, local GABAergic interneurons influenced the activity of liver-projecting parasympathetic cholinergic neurons in the DMV, thereby regulating the hepatic glucose output^48^. Norepinephrine-expressing neurons in the NTS controlled feeding behavior^49–51^. Interestingly, activation of the NE-expressing neurons in the NTS (NTS^NE^) projecting projections to CGRP-expressing neurons in the PBN pathway reduced^50^, whereas stimulation of the NTS^NE^ to agouti-related peptide (Agrp)-containing neurons in the arcuate nucleus of the hypothalamus pathway promoted, feeding^49^. The anti-obesity effect that we observed would be in part due to the fact that local interneurons and projection neurons in the NTS receives the inadequate information on excess energy balance via hepatic vagal sensory neurons. Consequently, this sensory impairment would cause an increase in energy expenditure instead of storing excess nutrients in the liver.

The decrease in hepatic steatosis we observed in mice without liver-projecting vagal sensory neurons is likely attributable to the decrease in DNL, which was accompanied by a reduction in fatty acid uptake in the liver. Our gene expression analysis revealed that the experimental mice exhibited a decrease in *Srebf1* and its target genes, including *Fasn* and *Scd1*. The significant reduction in *Cd36* expression observed in the livers of experimental mice using both qPCR and spatial transcriptomics is noteworthy. This decrease may have a significant impact on DNL since CD36 is involved not only in transporting fatty acids but also in SREBP1 processing in the liver^52^. Interestingly, HFD feeding appeared to have an opposite effect on the two transcription factors, *Irf6* and *Tbx3* expression in the PC of control mice compared to the experimental mice. In fact, *Irf6* overexpression in hepatocytes reduced hepatic steatosis^30^, while hepatic *Tbx3* deletion improved hepatic steatosis in mice fed a high-fat diet^31^. It is highly likely that these three transcription factors and *Cd36* could be crucial downstream targets of liver-projecting vagal sensory input. Furthermore, the enrichment analysis of DEGs between the two groups showed that each zonation possessed a unique GO term. These specific terms included “increased liver weight” in the PP group, “abnormal hepatocyte morphology” in the Mid group, and “abnormal circulating amino acid levels” in the PC group. This suggests that each zonation may have distinct impacts on the liver under obesogenic conditions, leading to different types of abnormalities.

The animal behavior tests we conducted showed that the experimental male mice were more inclined to explore novel and unprotected areas, suggesting that they were less anxious and more motivated than the control mice. Although it appears that impaired hepatic lipid metabolism significantly affects psychiatric disorders^13–15^, the underlying mechanisms remains poorly understood. Our results suggest that liver-innervating vagal sensory neurons may play an important role in developing psychiatric disorders in patients with obesity and MAFLD. Importantly, male mice lacking liver-innervating vagal sensory neurons demonstrated reduced anxiety-like behavior compared with male control mice, whereas females exhibited no difference in anxiety-like behavior. It is possible that liver-innervating vagal sensory neurons exhibit different responses to interoceptive signals, resulting in the distinct regulation of hepatic lipid metabolism and anxiety-like behavior between the sexes. An implantable vagus nerve stimulator has been approved by the Food and Drug Administration to treat epilepsy and depression. As we recently demonstrated that liver-innervating parasympathetic cholinergic neurons played a critical role in the development of hepatic steatosis in mice fed HFD^53^, modulating the liver-brain axis via the vagus nerve may offer a promising therapeutic approach for improving lipid metabolism, glucose homeostasis, and affective disorders in obesity and diabetes.

## Materials and Methods

### Animals

All mouse care and experimental procedures were approved by the Institutional Animal Care Research Advisory Committee of Albert Einstein College of Medicine.

Eight-nine-week-old Avil^CreERT2^ (stock # 032027) and Rosa26-floxed-STOP-Cas9-eGFP mice (stock# 024857) were purchased from the Jackson Laboratory. Mice were housed in cages at a controlled temperature (22 °C) with a 12:12 h light-dark cycle and fed a high-fat diet (HFD; Research Diets, D12492; 20% calories by carbohydrate, 20% by protein, and 60% by fat) for 10- 12 weeks, with water provided ad libitum. The mice were administered intraperitoneal injections of 75 mg tamoxifen/kg body weight on five consecutive days. The tamoxifen solution was prepared by dissolving it in a mixture of 90% corn oil and 10% ethanol

### Viral injection

We used AAVrg-Cre (Addgene 55636-AAVrg, titer; 2.2 x10^13^ pfu/ml), AAVrg-FLEX-GFP (Addgene 51502-AAVrg, titer; 2.3 x10^13^ pfu/ml), AAVrg-FLEX-jGCamp7s (Addgene 104491, titer; 2.5 x10^13^ pfu/ml). Avil^CreERT2^ mice received a total volume of 20 μl (4 μl per site/ 5 different sites) of viral injection into the medial and left lobes of the livers. A Hamilton syringe was used to inject over 40 min. The needle (30 G) was left for an additional 10 min to allow diffusion of the virus within the parenchyma. Mice were killed 5 and 10 weeks post-viral inoculation.

To ablate liver-innervating vagal sensory neurons, a total volume of 20 ul of AAVrg-FLEX-taCasp3-TEVp (titer >1 × 10^12^ pfu/ml; Addgene 45580) was injected into the medial and left lobes of the livers of Avil^CreERT2^ mice. AAVrg-FLEX-GFP (Addgene 51502-AAVrg, titer > 1 × 10^13^ pfu/ml) was used as control.

### Clear, unobstructed brain imaging cocktails (CUBIC) tissue clearing and immunostaining

The CUBIC method^54^ was used to clear brainstem tissues. Mice were perfused with 4% (w/v) paraformaldehyde (PFA) in PBS. Brain tissues were collected and post-fixed in 4% PFA at 4C° for 24 h. The tissue samples were sectioned using a vibratome at a thickness of 300 um. The sections were immersed in CUBIC reagent 1 (25 wt% urea, 25 wt% quadrol, and 15 wt% Triton X-100) with gentle shaking at 37C° for 3 days. After decolorization, the sections were washed three times with PBS at RT for 2 h and then incubated in a blocking solution composed of 1% donkey serum, 0.5% BSA, 0.3% Triton X-100, and 0.05% sodium azide for 3 h at room temperature (RT). Tissue samples were incubated with a rabbit anti-GFP (Rockland, 600-401- 215, 1:200) for 5 days at RT with mild shaking and washed three times with PBS (each for at least 2 h). Following wash-out, the tissue samples were incubated with an Alexa Fluor 488 donkey anti-rabbit IgG (1:200, Invitrogen, A21206) for 3 days at RT. The tissues were washed and immersed in CUBIC-reagent 2 (50 wt% sucrose, 25 wt% Urea, 10wt% Triethanolamine, and 0.1%(v/v) Triton X-100) for 1 d and mounted with a mixture of mineral oil and silicone oil. Z-stack images were obtained using a Leica SP8 confocal microscope, and 3D construction was performed using the AIVIA (version 12.1) and Leica application suite X.

### Immunostaining

Mice were anesthetized with isoflurane (3%) and transcardially perfused with 4% paraformaldehyde. Nodose ganglia, liver, and brain were post-fixed in 4% paraformaldehyde overnight in a cold room and then in 30% sucrose the following day. Tissues were sectioned using a cryostat at 16-20 μm. The sections were incubated with 0.3% Triton X-100 at room temperature (RT) for 30min, and then blocked in PBS buffer containing 5% donkey serum, 2% bovine serum albumin and 0.15% Triton X-100 for 1hr at RT and then incubated with goat anti-GFP (Novus, NB100-1770, 1:500), rabbit anti-advillin (Abcam, Ab72210, 1:1000), mouse anti-c-Myc (Invitrogen, 13-2500, 1:1,000) antibodies for overnight at RT, and then sections were washed three times in PBS and incubated with AlexaFluor 594 donkey anti-rabbit IgG (Jackson immunoresearch, 711-585-152, 1:500), AlexaFluor 488 donkey anti-goat IgG (Invitrogen, A11055, 1:500), and AlexaFluor 568 donkey anti-mouse IgG (Invitrogen, A10037, 1:500) for 1hr at RT. The sections were washed, dried, and mounted using VECTASHIELD medium containing DAPI. Images were acquired using a Leica SP8 confocal microscope.

For histological analysis, the liver tissues were embedded in paraffin and cut into 5 μm thickness. The sections were stained with hematoxylin and eosin. Lipid droplet accumulation was visualized by Oil Red O staining of the frozen liver sections. H&E and Oil Red O slides of liver sections were scanned using the Panoramic 250 FLASH III F2, and analyzed using slideViewer 2.7 (3DHISTECH).

To visualize vagal sensory nerve fibers in the liver, Avil^CreERT2^ mice were given an injection of AAVrg-CMV-FLEX-PLAP (with a titer of 1.0×10^12^ GC/ml from ABM, Inc.) into the liver. After two months following viral infections, the mice were perfused with PBS and fixed with a 4% paraformaldehyde solution. The liver tissues were then sliced to a thickness of 30 μm using a vibratome and incubated in alkaline phosphatase (AP) buffer (0.1 M Tris HCl pH 9.5, 0.1 M NaCl, 50 mM MgCl2, 0.1% Tween20, and 5 mM levamisole) for one hour and thirty minutes at room temperature. The tissue was then visualized using the NBT/BCIP substrate solution (ThermoFisher Scientific, 34042) until the desired stain developed. The tissues were washed with distilled water and mounted with aqueous mounting media (Vector, H-5501). The stained samples were captured in brightfield scanning mode using the Panoramic 250 FLASH III F2, and analyzed using slideViewer 2.7 (3DHISTECH).

### Measurement of body weight, body composition, and blood glucose

Body weight was measured weekly at 9 a.m. Body composition for fat mass and fat-free mass was assessed using the EchoMRI system. Blood samples were collected from the mouse

### Assessment of glucose tolerance and insulin tolerance

For the GTT, the experimental and control mice were fasted for 15 h (6:00 pm–9:00 am) 10 weeks post-viral inoculation. Sterile glucose solution was administered intraperitoneally at a concentration of 2 g/kg (glucose/body weight) at time zero. Blood glucose levels were measured at 15, 30, 60, 90, and 120 min after glucose injection. Blood glucose levels were plotted against time after glucose injection, and the area under the curve (AUC) was calculated and compared between experimental and control groups. For the ITT, the mice were fasted for 4 h (9:00 am to 1:00 pm). Blood glucose levels were measured at 0, 15, 30, 60, 90, and 120 min after intraperitoneal injection of human insulin (1 U/kg; Santa Cruz Biotechnology). We immediately injected glucose (2 g/kg) if the mice appeared ill due to insulin-induced hypoglycemia.

### Assessment of energy expenditure

Indirect calorimetry was performed for three days at the end of 10 weeks. O_2_ consumption and CO_2_ production were measured for each mouse at 10-min intervals over a 24-h period. Mice were individually housed in calorimeter cages (Oxymax, Columbus Instruments) and acclimated to the respiratory chambers for two days prior to gas exchange measurements. The respiratory exchange ratio was calculated as the ratio of CO_2_ production to the O_2_ consumption. Since there was no difference in food intake, VO_2_ was normalized to the lean body mass. Locomotor activity in the X-Y and Z planes was measured using infrared beam breaks in calorimetry cages.

### RT-qPCR

Liver tissues were collected from the experimental and control groups–10-12 weeks post viral inoculation. Liver tissues were homogenized in TRIzol reagent (ThermoFisher Scientific, 15596-018), and total RNAs were isolated according to the manufacturer’s instructions. First-strand cDNAs were synthesized using the SuperScript III First-Strand synthesis kit (ThermoFisher Scientific, 18080-051). qPCR was performed in sealed 96-well plates with SYBR Green I master Mix (Applied Biosystems, A25742) using a Quant Studio 3 system (Applied Biosystems). qPCR reactions were prepared in a final volume of 20 ul containing 2 ul cDNAs, and 10 ul of SYBR Green master mix in the presence of primers at 0.5 uM. Beta-actin (*Actb*) was used as an internal control for quantification of each sample. Amplification was performed under the following conditions: denaturation at 95□°C for 30 s, followed by 40 cycles of denaturation at 95□°C for 30 s and annealing/extension at 60□°C for 1 min.

We examined the following genes: *Srebp1c* (NM_001358315.1), *Chrebp* (NM_021455.5), *Acly* (NM_134037.3)*, Acaca* (NM_133360.3)*, Fasn* (NM_007988.3)*, Scd1* (NM_009127.4)*, Cd36* (NM_007643.5), *Fatp1* (NM_011977.3), *Fatp2* (NM_011978.2), *Fatp3* (NM_011988.3) *and Fatp4* (NM_011989.4). All the primer sequences were validated with “Primer BLAST” primer design program to ensure specificity for the target gene. Melt curves and gel electrophoresis were analyzed to confirm the specificity of the PCR products. Relative gene expression was determined using a comparative method (2^-ΔΔCT^).

### Single-nucleus RNA sequencing (snRNA-Seq)

Nuclei isolation and single-nucleus RNA sequencing were performed by the Singulomics Corporation (Singulomics.com, Bronx, NY, USA). The mice were euthanized with an overdose of isoflurane (≥ 5 %). Thirty vagus nerve ganglia were collected from eight male and seven female Rosa26-eGFP^f^ mice injected with AAVrg-Cre into the liver four weeks after viral injection^17^. Immediately after tissue collection, 16 vagus nerve ganglia from males and 14 vagus nerve ganglia from females were flash-frozen. Individual tissue samples were divided into separate pools for males and females. The pooled samples were subsequently homogenized and lysed using Triton X-100 in RNase-free water to isolate the nuclei. For nuclear isolation, we followed the 10x genomics protocol CG000124 (Rev. F). Briefly, isolated cells were centrifuged at 400 g for 5 min at 4°C and the supernatant was removed without disturbing the cell pellet. Cells were completely suspended in 200 μl lysis buffer and lysed on ice for 1 min. We added 800 μl of nuclei wash and resuspension buffer containing 1% bovine serum albumin and 0.2 U/μl RNase inhibitor and centrifuged the nuclei at 500 g for 10 min at 4°C. The isolated nuclei were diluted to 700 nuclei/µl for standardized 10x capture. The 10x Genomics single cell protocol was immediately performed using 10x Genomics Chromium Next GEM 3’ Single Cell Reagent kits v3.1 (10x Genomics, Pleasanton, CA). Libraries were sequenced using the Illumina NovaSeq 6000 system (Illumina, San Diego, CA, USA).

The libraries were sequenced with ∼200 million PE150 reads per sample using Illumina NovaSeq. Raw sequencing reads were analyzed with the mouse reference genome (mm10) with the addition of the human growth hormone poly(A) (hgGH-poly(A)) transgene sequence found in AAVrg-Cre using Cell Ranger v7.1.0. Results containing this information have been submitted elsewhere. Introns were included in the analyses. To further clean the data, Debris Identification using the Expectation Maximization (DIEM) program ^55^ was used to filter the barcodes (from raw_feature_bc_matrix of each sample), keeping barcodes with debris scores < 0.5, number of features (genes) > 250, and UMI count > 1000.

Aggregation of the samples was performed with the cellranger aggr function, normalizing for the total number of confidently mapped reads across libraries. We then used the cellranger reanalyze function to analyze the samples with only the filtered, barcodes. The UMAP is generated by the cellranger reanalyze function, running at the default setting. Differential gene expression was determined using the 10x Genomics Loupe Browser (version 7). It should be noted that single-cell DE analysis carries a high risk of false-positive detection of differentially expressed genes. The significant gene test in the Loupe Cell Browser replicated the differential expression analysis in Cell Ranger.

### Spatial Transcriptomics

Livers were snap-frozen in liquid nitrogen and thawed briefly on ice, followed by embedding in a cryomold (Sakura, 4565) filled with Neg-50 (Epredia, 6502). The cryomold was then frozen in a container with pre-chilled 2-methylbutane on dry ice. The samples were cryosectioned (Cryostar NX70 Cryostat, −18C for blade, −18 °C for specimen head, set at 10mm thickness) and placed on a pre-chilled Visium Spatial Gene Expression Slide (10x Genomics, PN-2000233). The tissue sections were processed following 10x Genomics protocols (PN- 1000187, PN-1000215). Adaptors and sequencing primers were obtained from Illumina. The libraries were sequenced by Novogene using a NovaSeq 6000 (PE150) instrument, targeting 80,000 reads per spot. Fastq files were aligned against mouse mm10 reference and histology images were processed with Spaceranger (2.0). Pericentral, mid-zone, and periportal clusters were identified based on graph-based clusters and known zonation markers (*Gstm3*, *Glul*, *and Gulo* for pericentral, *Cyp2f2*, *and Cdh1* for periportal). Violin plots were generated using the Loupe Browser (7).

### Measurement of hepatic TG and plasma insulin levels

Liver samples were harvested from mice with or without liver-innervating sensory neurons. Tissue samples were weighed and homogenized in 1 mL of a diluted NP-40 solution to solubilize triglycerides. The samples were centrifuged at 16,000 × g to remove the insoluble material. The TG levels were measured using a TG assay kit (Cayman Chemical, 10010303).

Blood samples from the experimental and control groups were collected from the retroorbital plexus using heparinized capillary tubes. Whole blood was centrifuged at 12,000 × g for 10 min, and plasma was separated and stored at −20°C until use. Plasma insulin concentrations were determined using two-site sandwich ELISA kits (Mercodia, 10-1247-0).

### Animal behavior tests

For the open field exploration test, mice were placed in the center of a chamber measured 40 cm (length) x 40 cm (width) x 40 cm (height) and allowed to explore the chamber for 10 min freely. The center region was designated as 20 x 20 cm^2^. The EthoVision XT video tracking system (Nodulus) was used to record the sessions and analyze the behavior, movement, and activity of the animals. The chamber was wiped with 95% ethanol before the subsequent tests to remove any scent clues. The total distance traveled, and time spent in the center and outer zones of the chamber were measured.

To perform the elevated plus maze test, mice were placed in the central area of the maze consisting of four arms in the shape of a plus sign elevated approximately 1 m from the ground. Mice were allowed to explore the maze for 5 min freely. The session was recorded, and the number of arm entries and amount of time spent in the open and closed arms were analyzed using the EthoVision XT video tracking system.

To perform the light-dark box test, mice were placed in the box that consists of two chambers, one light and one dark. We measured the time spent in each compartment and crossing from one compartment to the other using the video tracking system.

### Statistics

All statistical results are presented as the mean ± SEM. Statistical analyses were performed using GraphPad Prism (v.10). Two-tailed *t*-tests were used to calculate the p-values for pair-wise comparisons. Time-course comparisons between groups were analyzed using a two-way repeated-measures analysis of variance (RM ANOVA) with Sidak’s correction for multiple comparisons. Data were considered significantly different when the probability value was less than 0.05.

## General

We thank Dr. Shun-Mei Liu and Licheng Wu for their technical assistance.

## Funding

This work was supported by the NIH (R01 AT011653 and R03 MH137614 to Y.-H.J, R01 DK092246 to Y.-H.J and G.J.S., P30 DK020541 to Y.-H.J, G.J.S., and J.E.P. and DK110063 to J.E.P.).

## Competing interests

The authors declare no conflict of interest.

## Data availability

Single nucleus RNA-sequencing data were deposited into the Gene Expression Omnibus (GEO) database under accession number GSE267231. The spatial transcriptomic dataset was also submitted to GEO and assigned the accession number GSE264570.

## Notes

### Competing Interest Statement

The authors have declared no competing interest.

### Summary of Updates

This version of the manuscript has been revised to update new results and figures.

