## Supplementary materials for "Liver-innervating vagal sensory neurons are indispensable for the development of hepatic steatosis and anxiety-like behavior in diet-induced obese mice"

### Supplementary materials and methods

Liver-innervating vagal sensory neurons play an indispensable role in the development of hepatic steatosis and anxiety-like behavior in mice fed a high-fat diet.

Jiyeon Hwang^1,2^, Sangbhin Lee^1,2^, Junichi Okada^1,2^, Li Liu^1,2^, Jeffrey E. Pessin^1,2,3^, Streamson C. Chua Jr.^1,2,4^, Gary J. Schwartz^1,2,4^, and Young-Hwan Jo^1,2,3,4*^

**Affiliations**

^1^The Fleischer Institute for Diabetes and Metabolism

^2^Division of Endocrinology, Department of Medicine

^3^Department of Molecular Pharmacology,

^4^Department of Neuroscience,

Albert Einstein College of Medicine, NY, USA

For histological analysis, the liver tissues were embedded in paraffin and cut into 5 μm thickness. The sections were stained with hematoxylin and eosin. Lipid droplet accumulation was visualized by Oil Red O staining of the frozen liver sections. H&E and Oil Red O slides of liver sections were scanned using a fast scanner. Representative tissue areas were captured using Slideviewer software (version 2.6).

**Measurement of body weight, body composition, and blood glucose**

Body weight was measured weekly at 9 a.m. Body composition for fat mass and fat-free mass was assessed using the EchoMRI system. Blood samples were collected from the mouse tail, and a small drop of blood was placed on the test strip of a glucose meter.

**Single-nucleus RNA sequencing (snRNA-Seq)**

Nuclei isolation and single-nucleus RNA sequencing were performed by the Singulomics Corporation (Singulomics.com, Bronx, NY, USA). Mice were euthanized with overdose of isoflurane (5% or greater). Thirty vagus nerve ganglia were collected from eight male and seven female Rosa26-eGFP^f^  mice injected with AAVrg-Cre into the liver four weeks after viral injection ^2^. The transcriptome data of these GFP-positive liver-projecting vagal sensory neurons were described in the separate study ^2^. Immediately after tissue collection, 16 vagus nerve ganglia from males and 14 vagus nerve ganglia from females were flash-frozen. Individual tissue samples were divided into separate pools for males and females. These pooled samples were subsequently homogenized and lysed using Triton X-100 in RNase-free water to isolate the nuclei. For nuclear isolation, we followed the 10x genomics protocol, CG000124 (Rev. F). Briefly, isolated cells were centrifuged at 400 g for 5 min at 4°C and the supernatant were removed without disturbing the cell pellet. Cells were completely suspended in 200 μl lysis buffer and lysed on ice for 1 min. We added 800 μl nuclei wash and resuspension buffer containing 1% bovine serum albumin and 0.2 U/μl RNase inhibitor and centrifuged the nuclei at 500 g for 10 min at 4°C. The isolated nuclei were diluted to 700 nuclei/µl for standardized 10x capture. The 10x Genomics single cell protocol was immediately proceeded using 10x Genomics Chromium Next GEM 3' Single Cell Reagent kits v3.1 (10x Genomics, Pleasanton, CA). Libraries were sequenced using an Illumina NovaSeq 6000 (Illumina, San Diego, CA, USA).

To perform the light-dark box test, mice were placed in the box that consists of two chambers, one light and one dark. We measured time spent in each compartment and crossings from one compartment to the other with the video tracking system.

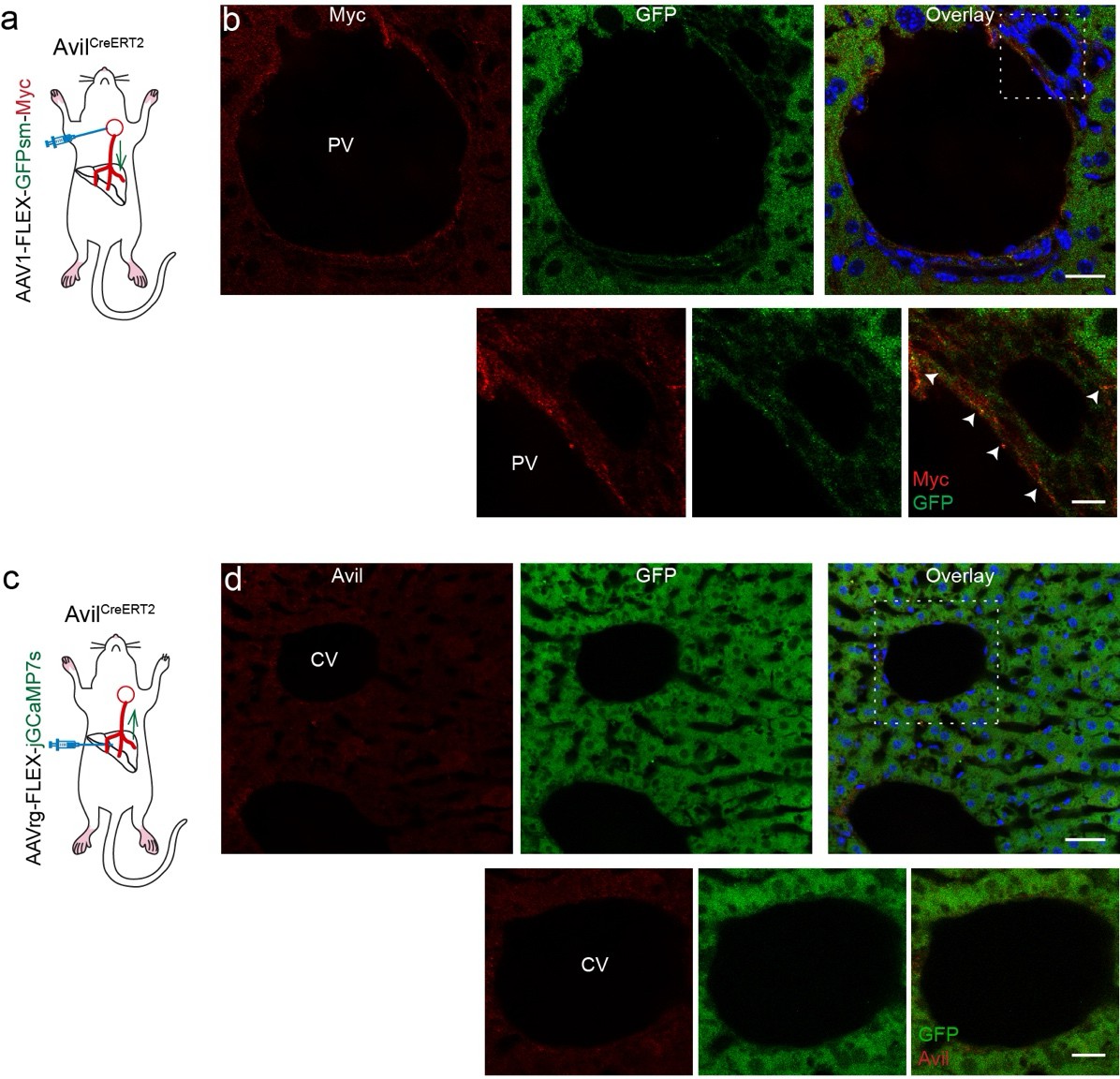

**Supplementary Figure 1**. Avil-positive vagal sensory neurons send axonal projections to the periportal area.

**a** and **b**. Schematic illustration depicting our experimental conditions. Anterograde monosynaptic AAV-FLEX-GFPsm-myc viruses were injected into the left nodose ganglion of of Avil^CreERT2^ mice (a). GFP- and Myc-positive nerve terminals were found in the portal vein area. Scale bar, 20 µm. Bottom panel: higher magnification view of the white square area. Arrowheads represent co-expression of Myc and GFP. Bottom panel: scale bar, 10 µm

**c** and **d**. Schematic illustration of the experimental conditions. AAVrg-FLEX-jGCaMP7s viruses were injected into the medial and left lobes of the livers of Avil^CreERT2^ mice. Images showing that there were no Avil (red)- and GFP-nerve terminals in the central vein area (d). Scale bar, 20 µm Bottom panel: higher magnification view of the white square area. Scale bar, 10 µm

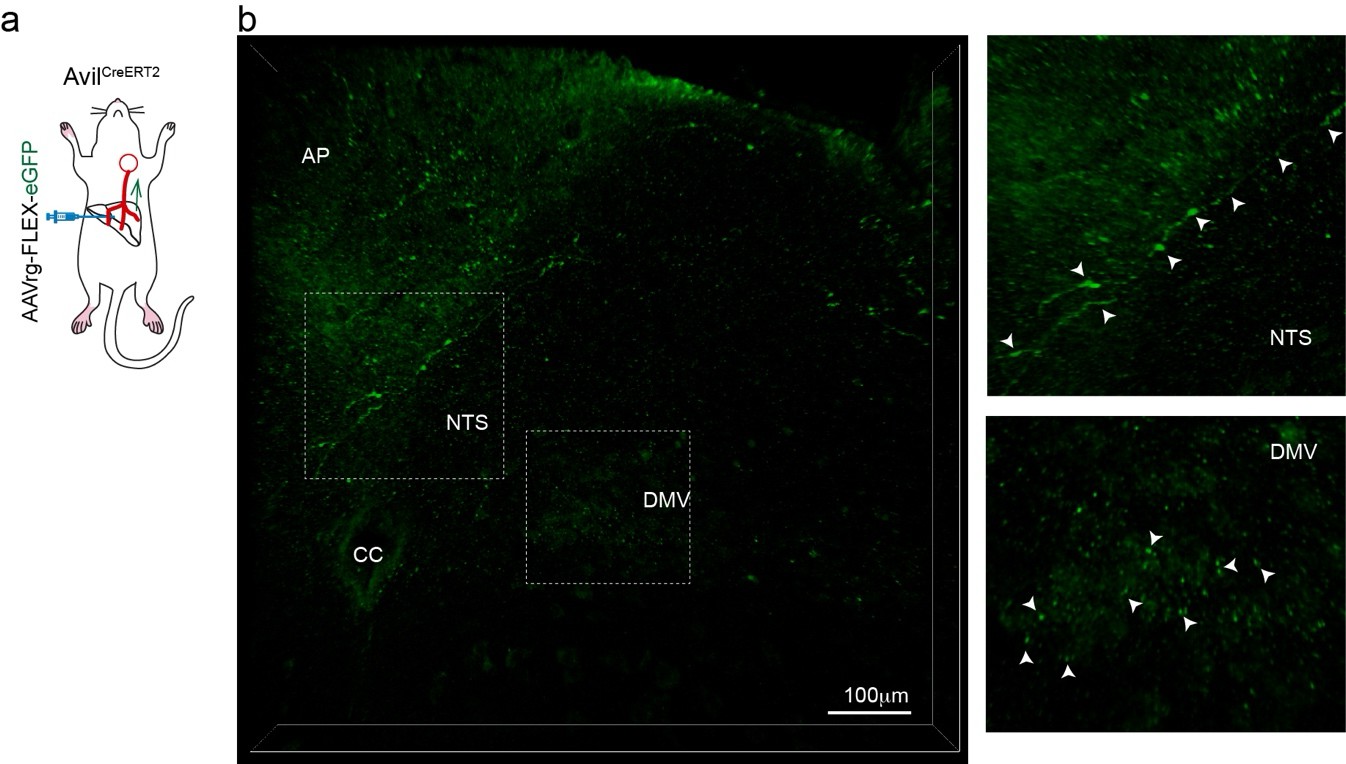

**Supplementary Figure 2**. Liver-innervating vagal sensory neurons project to the NTS, AP, and the DMV.

a. Schematic illustration of the experimental conditions. AAVrg-FLEX-eGFP viruses were injected into the medial and left lobes of the livers of Avil^CreERT2^ mice.
b. Z-stack projection images of confocal fluorescence microscopy showing GFP-positive nerve terminals in cleared brainstem tissue (thickness, 30 µm). Scale bar, 100 µm Right panels: higher magnification view of the white square area. Arrowheads represent GFP-positive axon terminals and puncta.

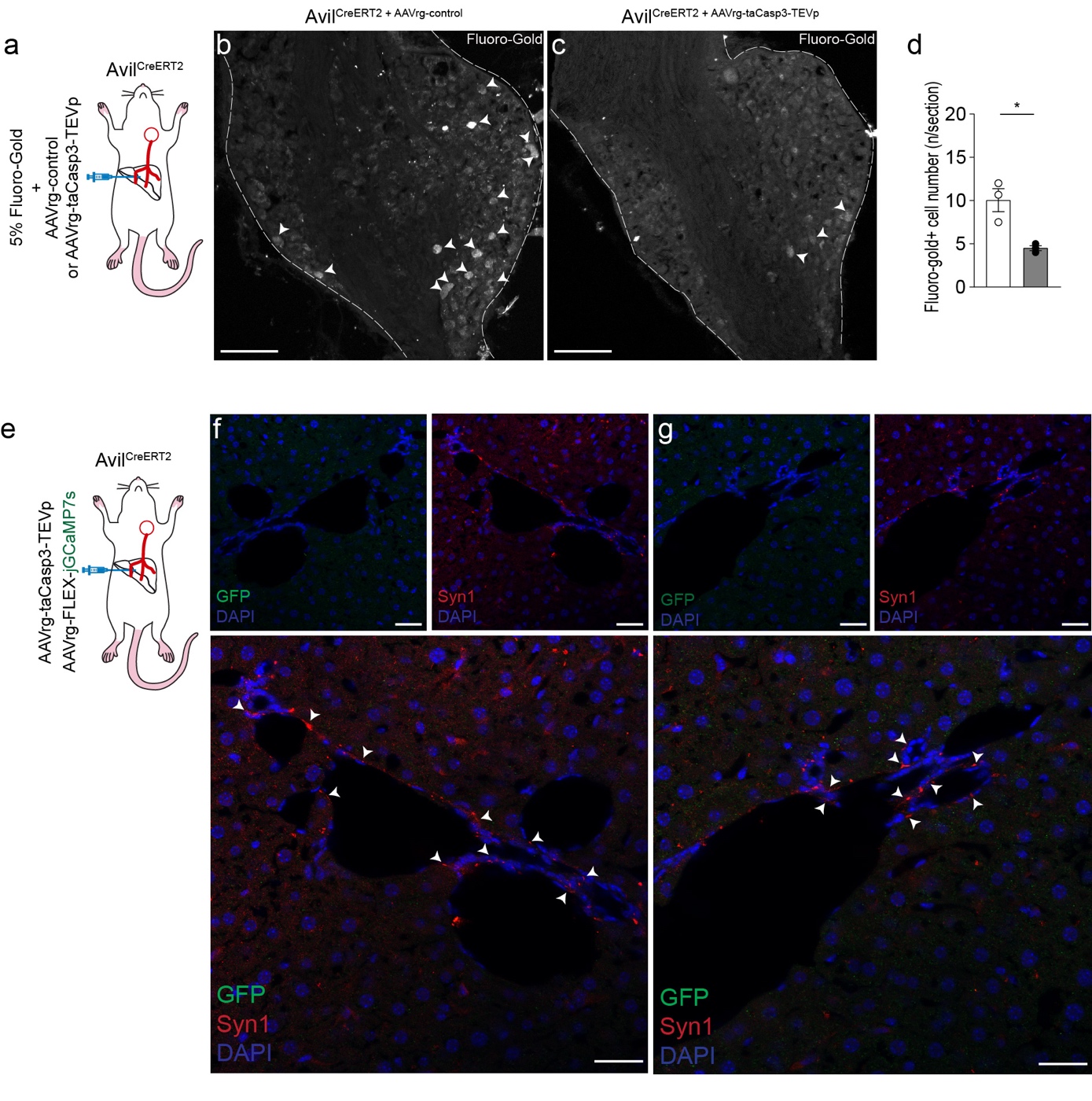

**Supplementary Figure 3**. Deletion of liver-projecting vagal sensory innervation in Avil^CreERT2^ mice injected with AAVrg-taCasp3-TEVp.

a. Schematic illustration of the experimental conditions. 5% of Fluoro-Gold was administered into the medial and left lobes of the liver of control and experimental mice.

**b** and **c**. Images showing Fluoro-Gold-positive vagal sensory neurons in the nodose ganglion in control and experimental mice. Scale bar, 100 µm

**d**. Graph showing Fluoro-Gold-positive vagal sensory neurons in the nodose ganglion in the control (n=3 mice) and experimental mice (n=3). Unpaired *t*-test, p<0.05

**e**. Schematic illustration of our experimental conditions. AAVrg-FLEX-jGCaMP7s viruses were injected to the liver of the experimental mice.

f and g. Images showing the presence of Syn1-positive nerve terminals, but not GFP-positive nerve terminals in the periportal areas. Bottom panel: Higher magnification view of the upper panel. Arrowheads represent Syn1-positive axon terminals and puncta. Scale bar, 20 µm

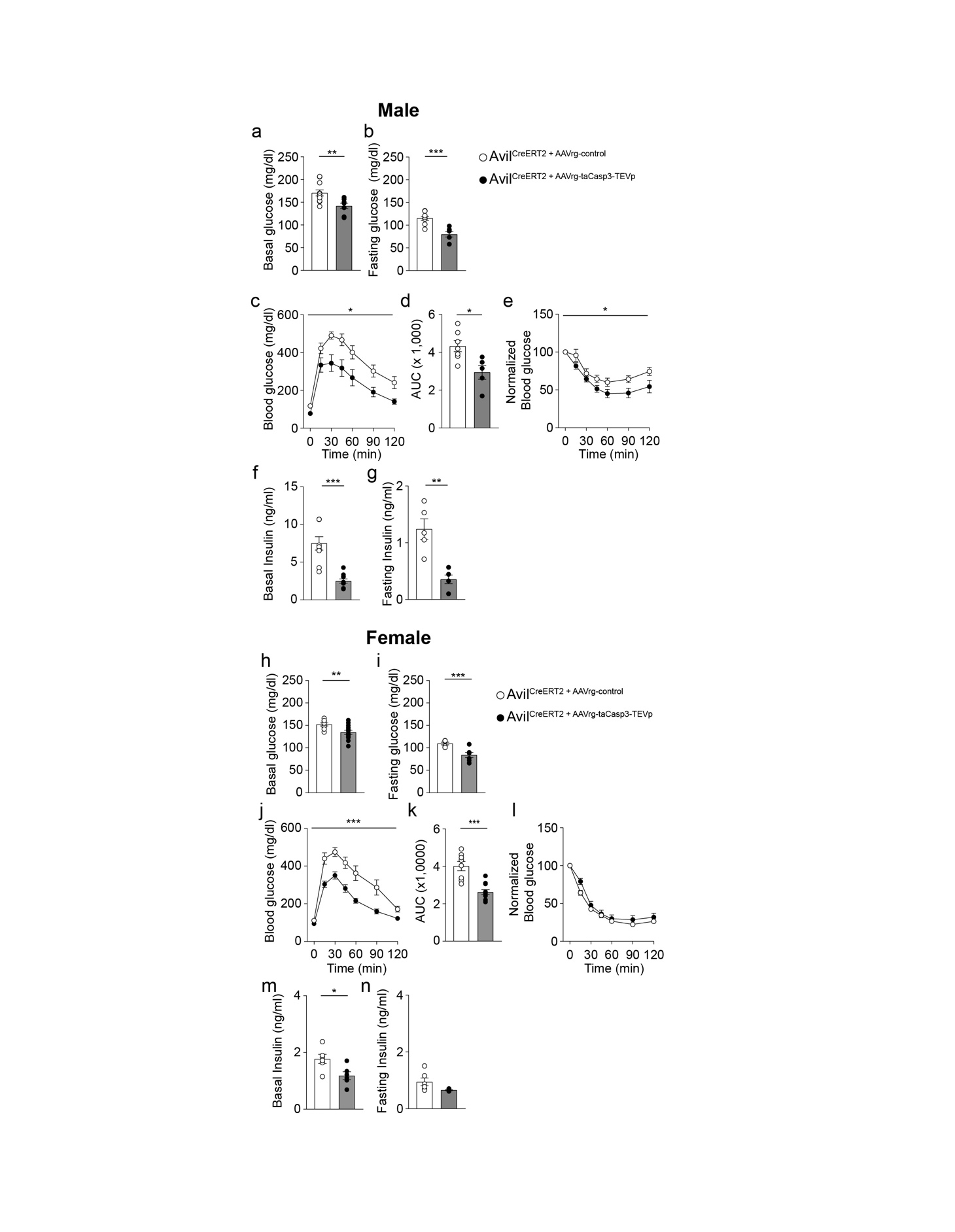

### Supplementary Figure 4. Deletion of liver-projecting vagal sensory innervation improves glucose tolerance and insulin sensitivity in male and female mice.

a. Graph showing that the basal glucose levels were significantly lower in the experimental groups than in the control group (control, n=11 mice; experimental, n=8 mice; unpaired *t*-test,

**p<0.01).

b. Graph showing the fasting glucose levels in the control and experimental groups (control, n= 9 mice; experimental, n= 6 mice; unpaired *t*-test, ***p<0.001).

**c** and **d**. Summary graphs showing the GTT in Avil^CreERT2^ mice with (n= 7 mice) and without (n= 5 mice) liver-projecting vagal sensory neurons (Two-way ANOVA followed by Sidak multiple comparison test, *p<0.05). AUC (**d**) of the control (n= 6 mice) and experimental (n = 5 mice; unpaired t-test, *p < 0.05) groups.

**e.** Plot showing the ITT in the experimental and control groups (control, n= 10 mice; experimental, n= 9 mice; two-way ANOVA followed by Sidak multiple comparisons test,

*p<0.05).

**f** and **g**. Graphs showing plasma basal and fasting insulin levels in the control (n= 9 and 5 mice, respectively) and experimental groups (n= 10 and 5 mice, respectively, unpaired *t*-test,

***p<0.001, **p<0.01).

**h** and **i.** Graphs showing basal (n= 9 vs. 13 mice) and fasting (n=7 vs. 13 mice) glucose levels in the control and experimental groups (unpaired *t*-test, **p<0.01, ***p<0.001).

**j** and **k**. Summary graphs showing the GTT in Avil^CreERT2^ female mice with (n= 8 mice) and without (n= 11 mice) liver-projecting vagal sensory neurons (two-way ANOVA followed by Sidak multiple comparison test, ***p<0.001). AUC (**k**, unpaired *t*-test, ***p<0.001).

**l.** Graph showing no difference in the ITT in Avil^CreERT2^ mice with and without liver-projecting vagal sensory neurons (control, n= 10 mice; experimental group, n= 14 mice).

**m** and **n**. Graphs showing basal plasma and fasting insulin levels in the control (n= 6 mice) and experimental groups (n= 6 mice, unpaired *t*-test, *p<0.05)

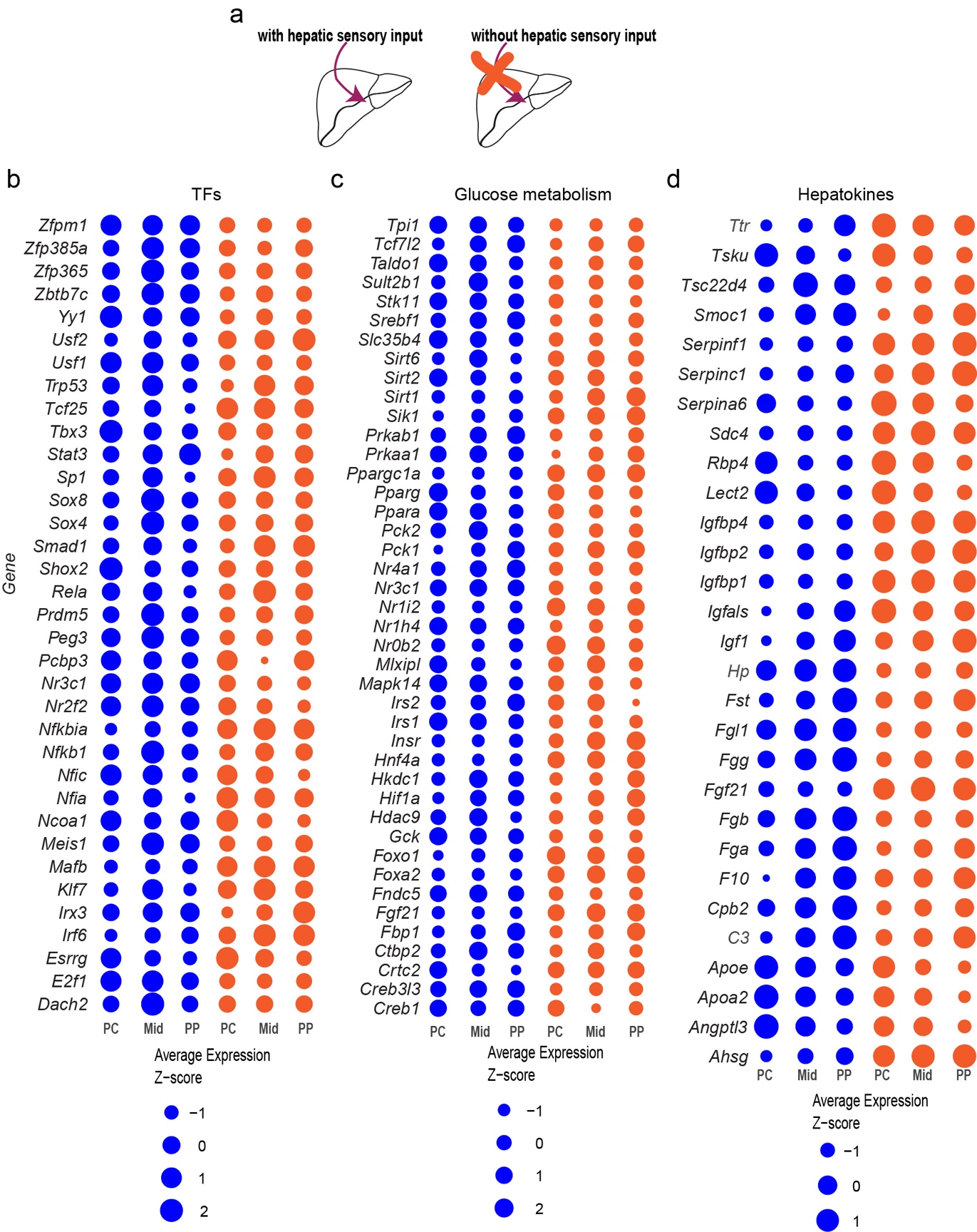

### Supplementary Figure 5. Differentially expressed genes identified in mouse livers with and without hepatic sensory innervation.

**a**. Schematic drawing of our experimental conditions.

**b**, **c**, and **d**. Bubble heatmaps displaying genes with differential expression for transcription factors, glucose metabolism, and hepatokines in mouse livers, comparing those with and without hepatic vagal sensory neurons.

**Supplementary Movie 1**. Z-stack projection images of confocal fluorescence microscopy showing GFP-positive nerve terminals in cleared brainstem tissue of Avil^CreERT2^ mice injected with AAVrg-FLEX-eGFP viruses into the medial and left lobes of the livers. GFP-positive axon terminals were detected in the NTS, AP, and DMV.

**Supplementary Table 1**. Differential gene expression in individual liver-projecting vagal sensory neurons (n= 6 neurons).

**Supplementary Table 2**. Differential gene expression in the pericentral zones of control and experimental mice.

**Supplementary Table 3**. Differential gene expression in the periportal zones of control and experimental mice.

**Supplementary Table 4**. Differential gene expression in the midlobular zones of control and experimental mice.
